# Neuronal nuclear calcium signaling suppression of microglial reactivity is mediated by osteoprotegerin after traumatic brain injury

**DOI:** 10.1101/2022.04.03.486874

**Authors:** Albrecht Fröhlich, Florian olde Heuvel, Rida Rehman, Sruthi Sankari Krishnamurthy, Shun Li, Zhenghui Li, David Bayer, Alison Conquest, Anna M. Hagenston, Albert Ludolph, Markus Huber-Lang, Tobias Boeckers, Bernd Knöll, Maria Cristina Morganti-Kossmann, Hilmar Bading, Francesco Roselli

## Abstract

**Background:** Traumatic Brain Injury (TBI) is characterized by massive changes in neuronal excitation, from acute excitotoxicity to chronic hyper- or hypoexcitability. Nuclear calcium signaling pathways are involved in translating changes in synaptic inputs and neuronal activity into discrete transcriptional programs which not only affect neuronal survival and synaptic integrity, but also the crosstalk between neurons and glial cells. Here we report the effects of blunting neuronal nuclear calcium signals in the context of TBI.

**Methods:** We used AAV vectors to express the genetically-encoded and nuclear-targeted calcium buffer parvalbumin (PV.NLS.mCherry) or the calcium/calmodulin buffer CaMBP4.mCherry in neurons only. Upon TBI, the extent of neuroinflammation, neuronal death and synaptic loss were assessed by immunohistochemistry and targeted transcriptome analysis. Modulation of the overall level of neuronal activity was achieved by PSAM/PSEM chemogenetics targeted to parvalbumin interneurons. The functional impact of neuronal nuclear Calcium buffering in TBI was assessed by quantification of spontaneous whisking.

**Results:** Buffering neuronal nuclear calcium unexpectedly resulted in a massive and long-lasting increase in the recruitment of reactive microglia to the injury site, which was characterised by a disease-associated and phagocytic phenotypes. This effect was accompanied by a substantial surge in synaptic loss and significantly reduced whisking activity. Transcriptome analysis revealed a complex effect of TBI in the context of neuronal nuclear calcium buffering, with upregulation of complement factors, chemokines and interferon-response genes, as well as the downregulation of synaptic genes and epigenetic regulators compared to control conditions. Notably, nuclear calcium buffering led to a substantial loss in neuronal osteoprotegerin (OPG). Whereas stimulation of neuronal firing induced OPG expression. Viral re-expression of OPG resulted in decreased microglial recruitment and synaptic loss. OPG upregulation was also observed in the CSF of human TBI patients, underscoring its translational value.

**Conclusion:** Neuronal nuclear calcium signals regulate the degree of microglial recruitment and reactivity upon TBI via, among others, osteoprotegerin signals. Our findings support a model whereby neuronal activity altered after TBI exerts a powerful impact on the neuroinflammatory cascade, which in turn contributes to the overall loss of synapses and functional impairment.

## Introduction

Traumatic Brain Injury (TBI) is characterized by the unfolding of multiple pathogenic cascades involving all cellular components of the central nervous system (CNS), including neurons, astrocytes, microglia, and several subpopulations of infiltrating immune cells (Morganti-Kossmann et al., 2019). The phenotype of reactive and infiltrating cells is not only determined by the extent of injury, but also by their reciprocal interactions as well as their relationships with non-immune cells, including neurons (Szepesi et al., 2018; Ouali-Alami et al., 2020).

Neuronal excitation plays a pivotal role in shaping the vulnerability of neurons in trauma. In fact, hyperexcitation, due to the excessive stimulation of extrasynaptic-type glutamate receptors, is thought to substantially contribute to secondary neuronal loss (secondary injury) occurring minutes-to-hours after the primary lesion (Yan et al., 2020; Stover et al., 1999; Pohl et al., 1999). In the excitotoxic phase, cytoplasmic Ca^2+^ overload impairs mitochondrial metabolism (Robertson, 2004), drives the calpain-mediated disassembly of the cytoskeleton (McGinn et al., 2009) and suppresses transcriptional responses (Bading et al., 2017). Surprisingly, several hours-to-days following brain injury, electrophysiological recordings have shown hypoexcitation of cortical neurons (Johnstone et al., 2014; Allitt et al., 2016). Stimulation of neuronal activity at these timepoints by glutamatergic agonists (Pohl et al., 1999), chemogenetics (Chandrasekar et al., 2018; Chandrasekar et al., 2019) and GABA receptor blockers (Zhang et al., 2019) appears to deliver protective signals by decreasing neuronal vulnerability in a neuronal nuclear calcium signaling dependent manner. These findings are in keeping with the concept that synaptically-driven neuronal excitation is neuroprotective through the activation of multiple transcription factors (TF) including CREB, SRF and ATF3 (Zhang et al., 2009; Zhang et al., 2011; Förstner et al., 2018, Förstner and Knöll 2019).

Interestingly, alterations in neuronal excitation (hyper- or hypo-excitation) can have a significant impact on local microglia signaling, ultimately affecting their activation state (Umpierre et al., 2020; Shaw et al., 1994). Upon TBI, microglial cells rapidly migrate to the damaged area of the brain and adopt a reactive phenotype to perform removal of cell debris, and pruning of synapses, with either neurotoxic or neuroprotective outcomes (Morganti-Kossmann et al., 2019; Jassam et al., 2017; Russo et al., 2016). Importantly, both the increase and decrease of neuronal firing results in the substantial modulation of microglial morphology and signaling, leading to increased microglial migration and extension of their processes. Thus, it is conceivable that the extreme changes in neuronal excitation occurring in the minutes, hours and days following TBI, may have important immune-regulatory consequences on microglia morphology and function.

The extent and pattern of neuronal firing is translated into short-term plasticity and long-term transcriptional programs through changes in nuclear calcium (NC) signaling (Buchtal et al., 2012, Bading 2013). Upon neuronal depolarization (Yu et al., 2017), increased levels of cytoplasmic calcium (Ca^2+^) evoke the elevation of NC, which, in turn, activates nuclearly localized Ca^2+^/Calmodulin-dependent kinases including CamKIV as well as Ca^2+^-sensitive TF such as DREAM (Bading et al., 2013; Mozolewski et al., 2021). NC drives the transcription of a substantial number of genes associated with the preservation of structural integrity of neurons (Mauceri et al., 2015) and with anti-apoptotic responses (Depp et al., 2018; Ahlgren et al., 2014). Blunting NC has been previously achieved by overexpression of the high-affinity Ca^2+^-binding protein parvalbumin in the nucleus, through a strong nuclear localization Signal (PV.NLS construct; Pusl et al., 2002). Alternatively, suppression of NC signaling has also been accomplished with the sequestration of nuclear Ca^2+^/Calmodulin complex by an engineered Ca^2+^/CaM buffer (CaMBP4; Wang et al., 1995). Both PV.NLS and nuclear CaMBP4 have been shown to strongly downregulate transcriptional responses associated with synaptic activity and NC (Schlumm et al., 2013; Bas-Orth et al., 2017; Pruunsild et al., 2017) and to regulate the morphology of dendritic arborizations (Mauceri et al., 2015).

In the present study, we set out to investigate whether blunting NC prior to TBI would affect neuronal vulnerability. To our surprise, we found that suppressing neuronal NC triggers a massive build-up of reactive microglia with disease-associated-like phenotypes and local loss of synapses (Rehman et al., 2021; Keren-Shaul et al., 2017. Notably, the comparison of transcriptomes revealed a distinct loss of osteoprotegerin (OPG) upon NC blunting, whereas microglial accumulation and synaptic loss are reversed by the neuronal re-expression of OPG. Thus, OPG emerges as a new NC-dependent mediator of neuron-microglia interactions in the neuroinflammatory response following TBI.

## Results

### 1. Buffering nuclear calcium in neurons enhances the early accumulation of microglia upon

Since NC regulates neuroprotective transcriptional programs, we investigated whether buffering NC by PV.NLS interfered with neuronal survival after TBI. Firstly, since NC is critical for the phosphorylation of CREB (pCREB), we evaluated the target engagement of PV.NLS by assessing the levels of pCREB after TBI (Li et al., 2014; Bell et al., 2013; Hardingham et al., 1997). AAV9 encoding for hSyn::PV.NLS-mCherry (or an empty vector for control) was injected into the somatosensory cortex of adult mice, generating >90% infection efficiency (30 days after injection >90% of NeuN+ cells were mCherry+ (Supp. Fig. 1A-B). Thirty days after AAV9 injection, mice were randomized to undergo either mild TBI (NSS score 0-1) or sham surgery. There were a total of four experimental groups: 1. empty vector (control) sham (CS); 2. control TBI (CT); 3. PV.NLS sham (PS); 4. PV.NLS TBI (PT), all of which were sacrificed 3h after treatment. CT samples displayed a significant increase in neuronal pCREB compared to CS (Chandrasekar et al., 2019), but the expression of PV.NLS largely blunted the upregulation of pCREB (Fig. 1A-B). At this time point no neuronal loss was detected, irrespective of PV.NLS expression (Fig. 1C). Thus, PV.NLS largely prevented the neuronal pCREB rise after TBI.

**Figure 1.**
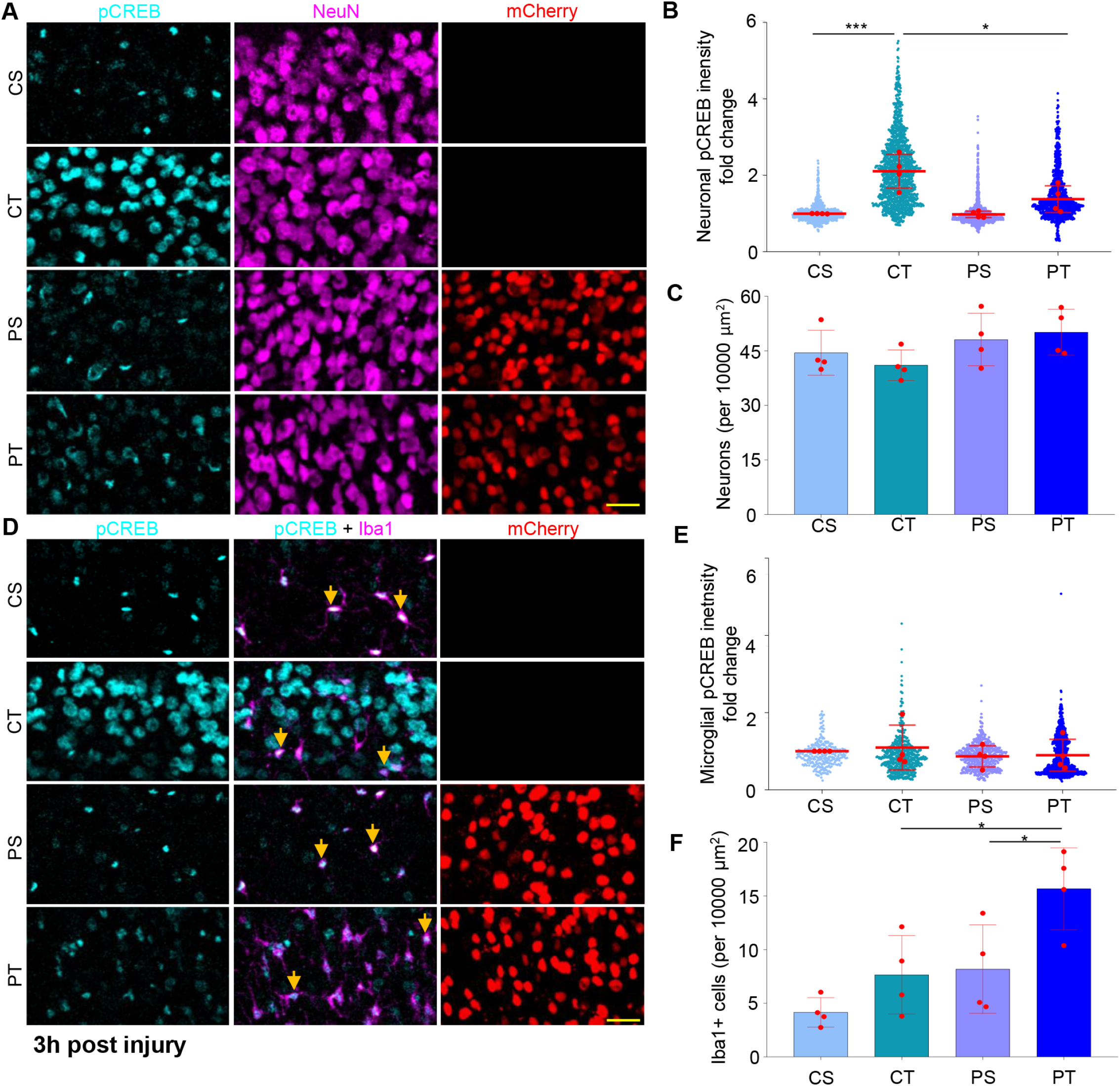
Buffering nuclear calcium in neurons enhances the accumulation of microglia 3h after TBI. A) Confocal images of the somatosensory cortex layer 2-3 obtained from Control Sham (CS), Control TBI (CT), PV.NLS Sham (PS) and PV.NLS TBI (PT) mice. Shown are pCREB (cyan), NeuN (magenta) and RFP (red). Scalebar: 25μm. B) Significant increase of neuronal pCREB intensity 3h post TBI compared to Sham (CS vs CT; 1.000 ± 0.004 vs 2.106 ± 0.441). Buffering of nuclear calcium significantly decreases neuronal pCREB intensity 3h post TBI (CT vs PT; 2.106 ± 0.441 vs 1.376 ± 0.350). Data is shown as a scatterplot with the mean of each animal indicated in red with mean ± SD. The mean of each animal was considered for statistical calculations. N = 4 mice. *: p< 0.05; ***: p < 0.001. C) No significant difference of neuronal density at the injury site 3h post TBI (CS vs CT; 44.460 ± 6.169 vs 41.050 ± 4.203) or post PV.NLS TBI (CT vs PT; 41.050 ± 4.203 vs 50.130 ± 6.344). Data are shown as mean ± SD. N = 4 mice. D) Confocal images of the somatosensory cortex layer 2-3 obtained from Control Sham (CS), Control TBI (CT), PV.NLS Sham (PS) and PV.NLS TBI (PT) mice. Shown are pCREB (cyan), IBA1 (magenta) and RFP (red). Arrows indicate double positive cells. Scalebar: 25 μm. E) No significant increase of microglial pCREB intensity 3h post TBI compared to sham (CS vs CT; 1.000 ± 0.000 vs 1.096 ± 0.581). Buffering of nuclear calcium does not significantly increase microglial pCREB intensity 3h post TBI (CT vs PT; 1.096 ± 0.581 vs 0.897 ± 0.411). Data is shown as a scatterplot with the mean of each animal indicated in red with mean ± SD. The mean of each animal was considered for statistical calculations. N = 4 mice. F) No significant difference in microglia density at the injury site 3h post TBI (CS vs CT; 4.156 ± 1.375 vs 7.651 ± 3.655). Buffering of nuclear calcium significantly increased microglia density post TBI (CT vs PT; 7.651 ± 3.655 vs 15.66 ± 3.812). Data is shown as mean ± SD. N = 4 mice. *: p < 0.05.

Analysis of brain sections immunostained for pCREB, showed a massive increase in a population of cells with small, elongated nuclei, highly immunopositive for pCREB in PT samples, which were barely detectable in CS and CT samples. Co-immunostaining of pCREB with GFAP and IBA1 revealed that almost all the pCREB+, small, elongated nuclei were detected in IBA1+ cells and therefore identified as microglia (>98%; (Supp. Fig. 1C-D)).

In an independent set of experiments, we verified that at 3h post-TBI, CT samples displayed only a small increase in IBA1+ cells compared to CS, whereas PT samples showed a massive increase in the number of IBA1+ cells (Fig. 1D, F). Furthermore, pCREB levels in microglia were not significantly altered in (Fig. 1D-E). Thus, buffering of neuronal NC in the acute phases of TBI unexpectedly resulted in the massive increase in local microglia.

### 2. Buffering of neuronal nuclear calcium induces a disease associated microglia (DAM)-like phenotype upon TBI

We characterized the IBA1+ population expanded in PT by immunostaining with the microglia marker TMEM119 and the disease-associated microglia (DAM)-like markers CD11c and CST7 (Rehman et al., 2021; Keren-Shaul et al. 2017). We also determined the expression of CD169, a marker of pathogenic phagocytes (Bogie et al., 2018; Rajan et al., 2020).

Across the four experimental groups, over >95% of IBA1+ cells were also TMEM119+ at 3h post injury (Fig. 2A, C), indicating that the contribution of infiltrating peripheral cells was comparatively minimal at this time point. Interestingly, PT samples showed the highest density of TMEM119+ cells (Fig.2B).

**Figure 2.**
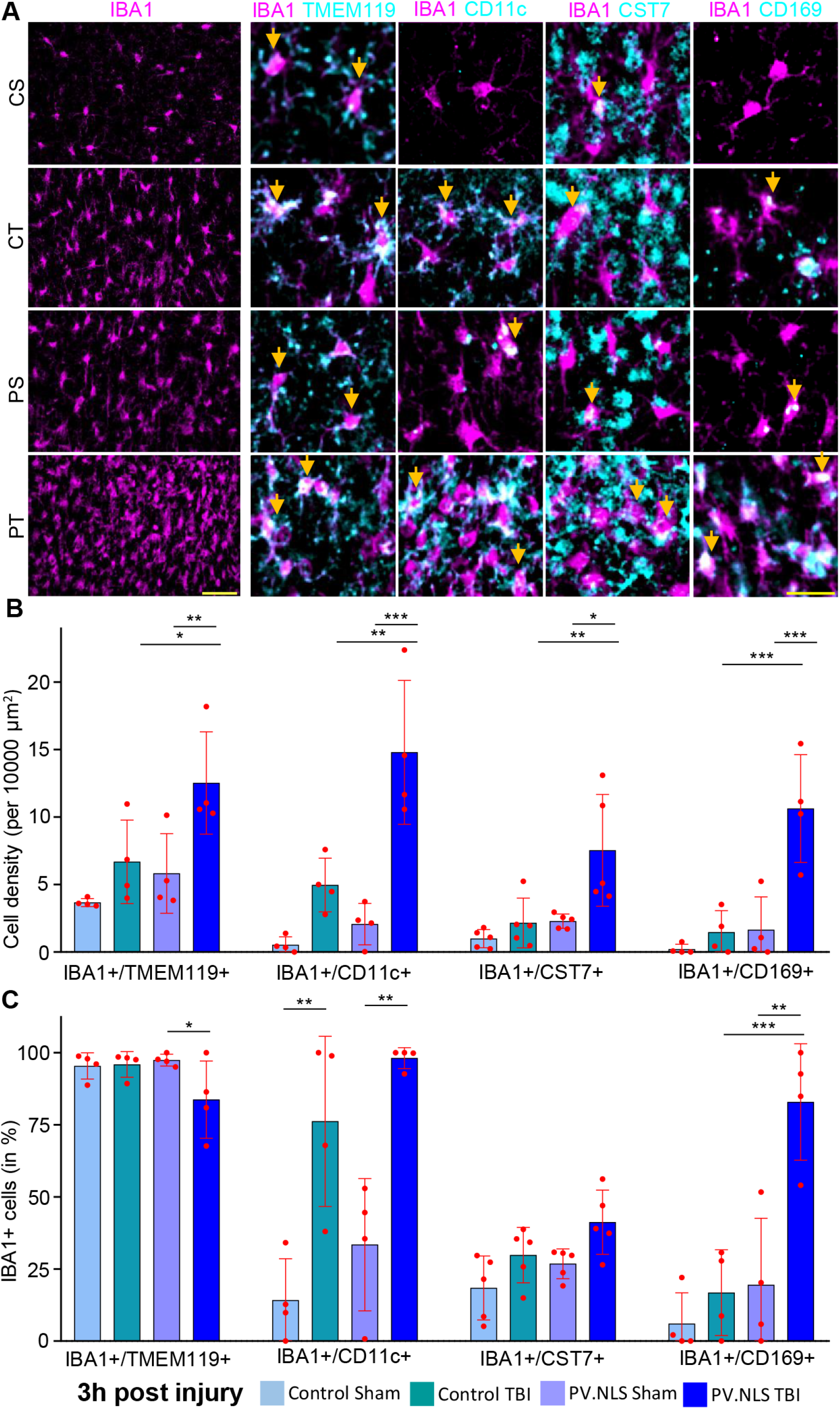
Buffering neuronal nuclear calcium induces a disease associated microglia (DAM)-like phenotype upon TBI. A) Confocal images of IBA1+ cells from the somatosensory cortex layer 2-3 obtained from Control Sham (CS), Control TBI (CT), PV.NLS Sham (PS) and PV.NLS TBI (PT) mice. In all the images IBA1 (magenta) is shown, additionally TMEM119, CD11c, CST7 and CD169 (all cyan) are shown. Arrows indicate double positive cells. Scale Bar 5Oμm (overview) and 25μm (inserts). B) Significant increase in IBA1+/TMEM119+ cells, IBA1+/CD11c+ cells and IBA1+/CST7+ cells 3h post TBI with nuclear calcium buffering compared to TBI alone (CT vs PT; for IBA1+/TMEM119+ cells 6.686 ± 3.090 vs 12.520 ± 3.791; for IBA1+/CD11c+ cells 4.964 ± 1.992 vs 14.800 ± 5.331; for IBA1+/CST7+ cells 2.151 ± 1.844 vs 7.533 ± 4.146). Significant increase in IBA1+/CD169+ cells 3h post TBI with nuclear calcium buffering compared to nuclear calcium buffering sham (PS vs PT; 1.638 ± 2.446 vs 10.630 ± 3.989). Data is shown as mean ± SD. N = 4-5 mice. *: p < 0.05; **: p < 0.01; *** p < 0.001. C) Significant decrease in percentage of IBA1+ cells copositive for TMEM119 3h post TBI with nuclear calcium buffering compared to TBI alone (CT vs PT; 95.92 ± 4.44% vs 83.75 ± 13.4%). Significant increase in percentage of IBA1+ cells copositive for CD11c 3h post TBI (CS vs CT; 14.18 ± 14.37% vs 76.19 ± 29.49%), but no effect of nuclear calcium buffering with TBI compared to TBI alone (CT vs PT; 76.19 ± 29.49% vs 98.13 ± 3.64%). Significant increase in percentage of IBA1+ cells copositive fro CD169 after nuclear calcium buffering and TBI compared to TBI alone (CT vs PT; 16.80 ± 14.86% vs 82.90 ± 20.20%). No significant effect observed in the percentage of IBA1+ cells copositive for CST7. Data is shown as mean ± SD. N = 4-5 mice. *: p < 0.05; **: p < 0.01; *** p < 0.001.

The density of IBA1+ CD11c+ cells was massively increased in PT samples compared to either CT or PS (Fig. 2A-B); however, their portion compared to overall IBA1+ population was unchanged (Fig. 2C). Likewise, the density of IBA1+/CST7+ cells was significantly increased in PT samples compared to CS, CT or PS but the presence of total IBA1+ cells was comparable in CT and PT. Taken together, these findings indicate that the increase in the number of DAM-like cells is due to an overall increase in microglial recruitment, but not to the enhanced commitment to a specific phenotype. On the other hand, only very few IBA1+ cells co-expressed CD169 in CS, CT or PT samples, but the number of IBA1+/CD169+ cells was massively increased in PT brains. Most notably, the population of IBA1+/CD169+ cells was also substantially increased in PT, implying that the strong expression of CD169+ in microglia corresponded to a reactive phenotype following the NC blockade as well as TBI (Fig. 2A, C). Thus, after TBI, the blockade of NC results in the appearance of a large microglial population with a DAM-like and unique phenotype characterized by high CD169 expression.

### 3. Blockade of neuronal nuclear calcium/calmodulin signaling recapitulates the enhanced recruitment of microglia after TBI

Since Ca^2+^/CaM-dependent kinases play a significant role in regulating signaling downstream of NC (Matthews et al. 1994; Schlumm et al. 2013), we hypothesized that sequestering nuclear Ca^2+^/CaM would recapitulate the effect of blunting NC with PV.NLS. For this purpose, we expressed a CaMBP4.mCherry construct, designed to bind and sequester nuclear Ca^2+^/CaM (Wang et al., 1995) or an empty vector for control in neurons of the somatosensory cortex via AAV injection. We considered four experimental groups: 1. control sham (CS); 2. control TBI (CT); 3. CaMBP4 sham (CaS) and 4. CaMBP4 TBI (CaT). Following TBI, the expression of CaMBP4 significantly reduced the upregulation of pCREB (Fig. 3A-B) but did not affect the vulnerability of neurons 3h after TBI (Fig 3C). Likewise, the expression of neuronal CaMBP4 caused a massive increase in microglial density upon TBI (Fig. 3D, F), but was ineffective in sham treated animals. Finally, we found that, compared to CS, the levels of pCREB were substantially increased in microglial cells when CaMBP4 was expressed in neurons (Fig. 3D, E). Thus, the inhibition of CaM-dependent NC signaling exerted by CaMBP4 in neurons largely recapitulates the effects of NC buffering by PV.NLS. This data suggests that NC-dependent action on CREB phosphorylation and microglial accumulation is specific and mediated by a CaM-dependent pathway.

**Figure 3.**
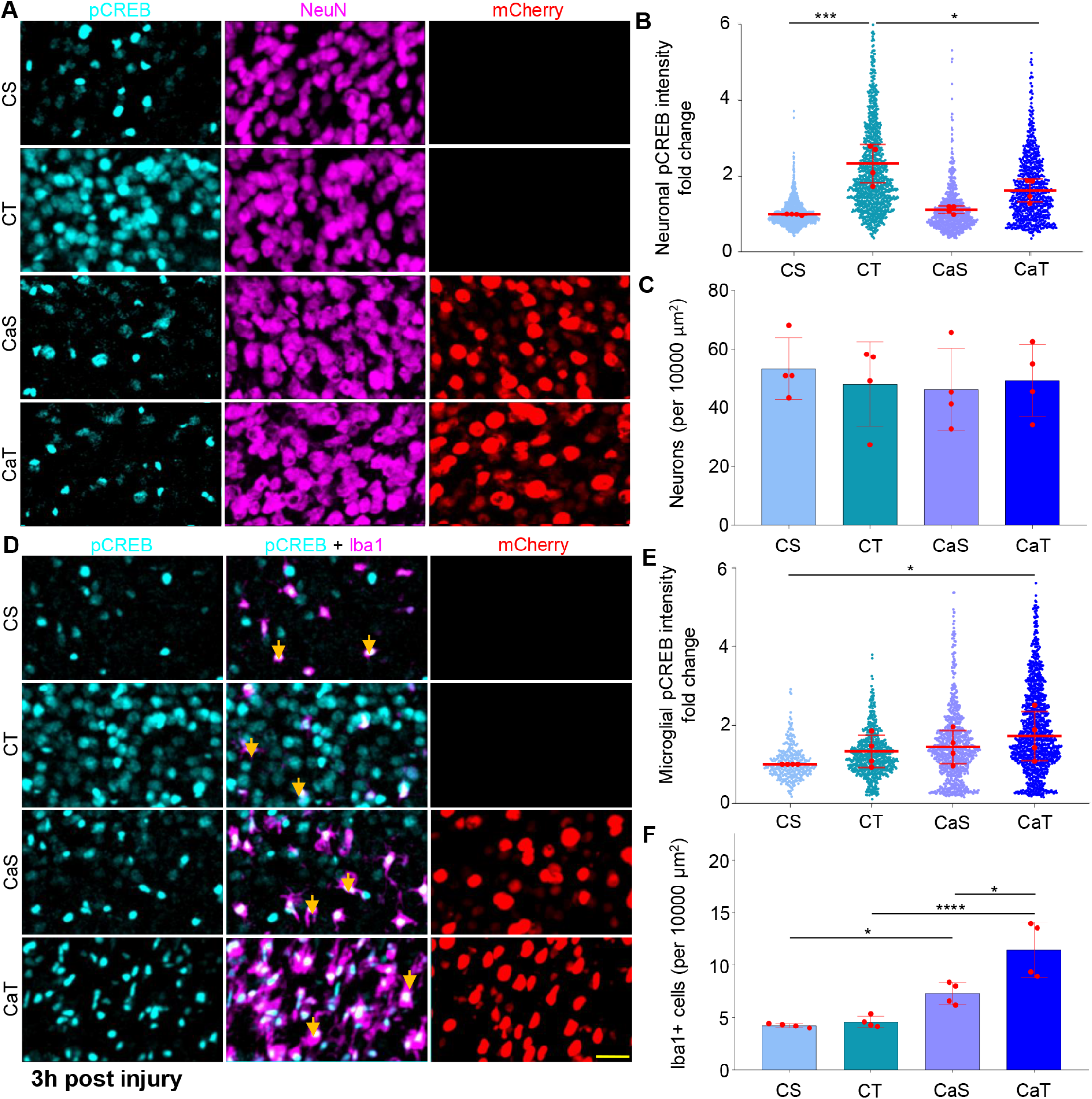
Blockade of neuronal nuclear calcium/calmodulin pathway recapitulates the enhanced recruitment of microglia after trauma. A) Confocal images of the somatosensory cortex layer 2-3 obtained from Control Sham (CS), Control TBI (CT), Calmodulin Sham (CaS) and Calmodulin TBI (CaT) mice. Shown are pCREB (cyan), NeuN (magenta) and mCherry (red). Scalebar: 25μm. B) Significant increase of neuronal pCREB intensity 3h post TBI compared to sham (CS vs CT; 1.000 ± 0.019 vs 2.332 ± 0.505). Buffering of CamK activation significantly decreases neuronal pCREB intensity 3h post TBI (CT vs CaT; 2.332 ± 0.505 vs 1.625 ± 0.299). Data is shown as a scatterplot with the mean of each animal indicated in red with mean ± SD. The mean of each animal was considered for statistical calculations. N = 4 mice. *: p < 0.05; ***: p < 0.001. C) No significant difference of neuronal density at the injury site 3h post TBI (CS vs CT; 53.51 ± 10.45 vs 48.04 ± 14.36) or post CaMBP4 TBI (CT vs CaT; 48.04 ± 14.36 vs 49.28 ± 12.22). Data is shown as mean ± SD. N = 4 mice. D) Confocal images of the somatosensory cortex layer 2-3 obtained from Control Sham (CS), Control TBI (CT), Calmodulin Sham (CaS) and Calmodulin TBI (CaT) mice. Shown are pCREB (cyan), NeuN (magenta) and mCherry (red). Arrows indicate IBA1+ cells expressing pCREB. Scalebar: 25 μm. E) No Significant increase of microglial pCREB intensity 3h post TBI compared to sham (CS vs CT; 1 ± 0 vs 1.332 ± 0.4122). Buffering of CamK activation does not significantly increase microglial pCREB intensity 3h post TBI (CT vs CaT; 1.332 ± 0.4122 vs 1.724 ± 0.6205). Data is shown as a scatterplot with the mean of each animal indicated in red with mean ± SD. The mean of each animal was considered for statistical calculations. N = 4 mice. **** p < 0.0001. F) No significant difference in microglia density at the injury site 3h post TBI (CS vs CT; 4.238 ± 0.187 vs 4.587 ± 0.531). Buffering of CamK activation significantly increased microglia density post TBI (CT vs CaT; 4.587 ± 0.531 vs 11.440 ± 2.665). Data is shown as mean ± SD. N = 4 mice. *: p < 0.05; ****: p < 0.0001.

### 4. Buffering of neuronal nuclear calcium enhances subacute microgliosis and synapse loss in TBI

We then explored the impact of NC buffering in TBI observed at later stages such as 24h post injury (1dpi) and 7d post injury. In agreement with the spatially heterogeneous nature of TBI lesions at these time point, we considered two regions of interest: one located at the center of the lesion (“core”) and the other located at a fixed lateral distance from the core (“perilesional area”; Supp. Fig. 2A) (Chandrasekar et al., 2018; Chandrasekar et al., 2019).

At 24h post injury, CT brains displayed an increased density of IBA1+ cells compared to sham mice (CT vs CS). However, in PT samples there was no evidence for a larger expansion of the IBA1+ population compared to CT samples, whereby cells showed a distinct ameboid morphology (Fig. 4A-B). The abundant IBA1+ population was associated with a significantly higher density (cells/area unit) of IBA1+/TMEM119+ cells in the core and perilesional areas (Fig. 4A-B and Supp. Fig. 2B-C). In addition, virtually all IBA1+ cells were TMEM119+, indicating a limited contribution of peripheral immune cells within and around the lesion (Fig. 4C). The IBA1+/CD11c+ subpopulation was substantially larger in PT samples (Fig. 4A-B), yet maintaining a comparable component of total IBA1+ cells in PT and CT brains, suggesting an overall expansion of the microglial population in PT.

**Figure 4.**
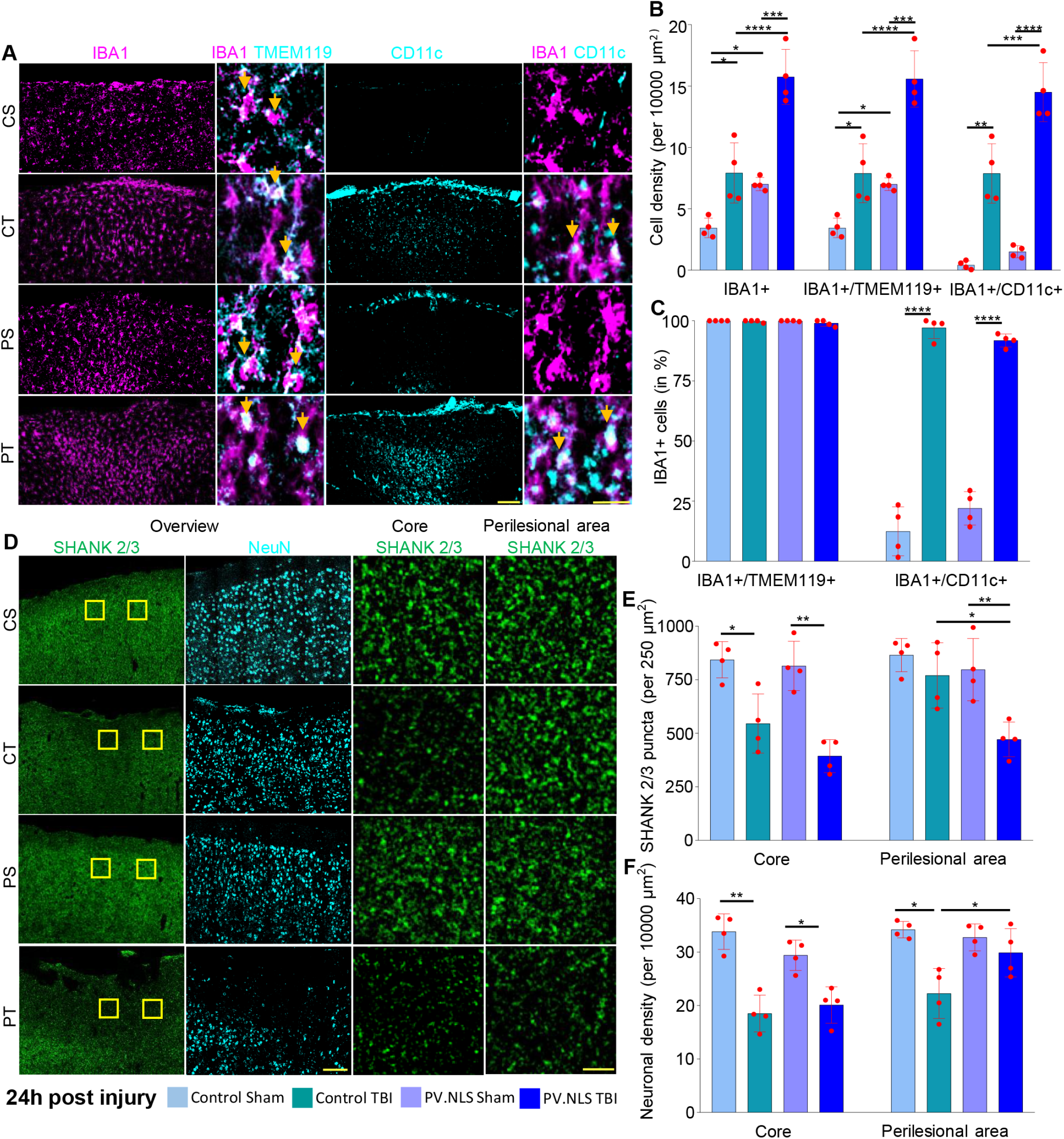
Buffering of neuronal nuclear calcium enhances subacute microgliosis and synapse loss 24h post trauma. A) Confocal images of the somatosensory cortex layer 2-3 obtained from Control Sham (CS), Control TBI (CT), PV.NLS Sham (PS) and PV.NLS TBI (PT). Shown are IBA1 (magenta), TMEM119 (cyan) and CD11c (cyan). Arrows indicate double positive cells. Scale bar 100μm (overview) and 20 μm (insert). B) Significant increase of IBA1+, IBA1+/TMEM119+ and IBA1+/CD11c+ cells 24h post TBI compared to sham (CS vs CT; for IBA1+ 3.440 ± 0.796 vs 7.917 ± 2.447; for IBA1+/TMEM119+ 3.44 ± 0.796 vs 7.893 ± 2.407; for IBA1+/CD11c+ 0.4167 ± 0.34 vs 7.881 ± 2.419). Buffering of nuclear calcium signaling in TBI significantly increased the density of IBA1+, IBA1+/TMEM119+ and IBA1+/CD11c+ cells (CT vs PT; for IBA1+ cells 7.917 ± 2.447 vs 15.740 ± 2.239; for IBA1+/TMEM119+ cells 7.893 ± 2.407 vs 15.570 ± 2.300; for IBA1+/CD11c+ cells 7.881 ± 2.419 vs 14.500 ± 2.403). Data is shown as mean ± SD. N = 4 mice. *: p < 0.05; **: p < 0.01; ***: p < 0.001; ****: p < 0.0001. C) No difference in percentage of IBA1+/TMEM119+ cells between treatment groups (CS 100.00 ± 0.00%; CT 99.75 ± 0.49%; PS 99.92 ± 0.17%; PT 98.98 ± 1.28%). Significant increase in percentage of IBA1+/CD11c+ cells 24h post TBI compared to sham (CS vs CT; 12.43 ± 10.19% vs 97.06 ± 4.48%). Buffering of nuclear calcium signaling in TBI did not alter the percentage of IBA1+/CD11c+ cells (CT vs PT; 97.06 ± 4.48% vs 91.8 ± 2.72%). Data is shown as mean ± SD. N = 4 mice. ***: p < 0.001. D) Confocal images of somatosensory cortex layer 2-3 obtained from Control Sham (CS), Control TBI (CT), PV.NLS Sham (PT) and PV.NLS TBI (PT). Shown are SHANK2/3 (green) and NeuN (cyan). Yellow inserts show analyzed core and perilesional areas. Scale bar 100μm (overview) and 5μm (insert). E) Significant decrease in core synaptic density 24h post TBI compared to sham (CS vs CT; 843.1 ± 84.6 vs 545.3 ± 138.2). Buffering of nuclear calcium signaling did not alter core synaptic density compared to TBI (CT vs PT; 545.3 ± 138.2 vs 393.2 ± 77.0). No difference in synaptic density 24h post TBI in the perilesional area compared to sham (CS vs CT; 864.9 ± 77.3 vs 769.9 ± 152.8). Buffering of nuclear calcium signaling significantly decreased synaptic density in the perilesional area compared to TBI (CT vs PT; 769.9 ± 152.8 vs 471.3 ± 80.9). Data is shown as mean ± SD. N = 4 mice. *: p < 0.05; **: p < 0.01. F) Significant decrease in core neuronal density 24h post TBI compared to sham (CS vs CT; 33.82 ± 3.32 vs 18.50 ± 3.44). Buffering nuclear calcium signaling did not alter neuronal density in the core compared to TBI alone (CT vs PT; 18.50 ± 3.44 vs 20.09 ± 3.41). Significant decrease in neuronal density in the perilesional area post TBI to sham and nuclear calcium buffering (CS vs CT; 34.2 ± 1.55 vs 22.25 ± 4.67; CT vs PT; 22.25 ± 4.67 vs 29.88 ± 4.52). Data is shown as mean ± SD. N=4 mice. *: p < 0.05; **: p < 0.01.

To assess the relationship between microglia infiltration with the extent of synapse loss, we quantified the integrity of the cortical architecture by measuring the density of excitatory synapses (number of pan-Shank+ puncta per area unit) and the number of surviving neurons in the core and perilesional brain areas.

In CT brains, a significant loss of synapses was observed in the core of the lesion, whereas in the perilesional area the synaptic density was only slightly reduced compared to CS brains (Fig. 4D-E). In contrast, we detected a robust loss of synapses in PT samples particularly in the perilesional area, indicating that the area of synaptic involvement is much larger than expected, as clearly manifested in the low-magnification pictures (Fig. 4D). Together, these experiments suggest that NC blockade resulted in a more extensive microgliosis and synaptic loss in the cortical area affected by TBI at 24h post injury. Neuronal density was substantially decreased in the core of both CT and PT groups (Fig. 4F). Interestingly, an increased preservation of neuronal density was found in the perilesional area of PT brains despite the increased microgliosis and synaptic loss.

At 7dpi, CT brains still displayed a slight accumulation of microglia, however their density remained substantially higher in PT sections both in the core and perilesional areas (Fig. 5A-B). Although there was little difference in neuronal loss between CT and PT both in the core and in the perilesional area (Fig. 5A, C), PT brains still displayed substantially reduced counts of synapses in the perilesional area (Fig. 5D-F), indicating the persistent larger area of synaptic loss which was already established at 24h post injury.

**Figure 5.**
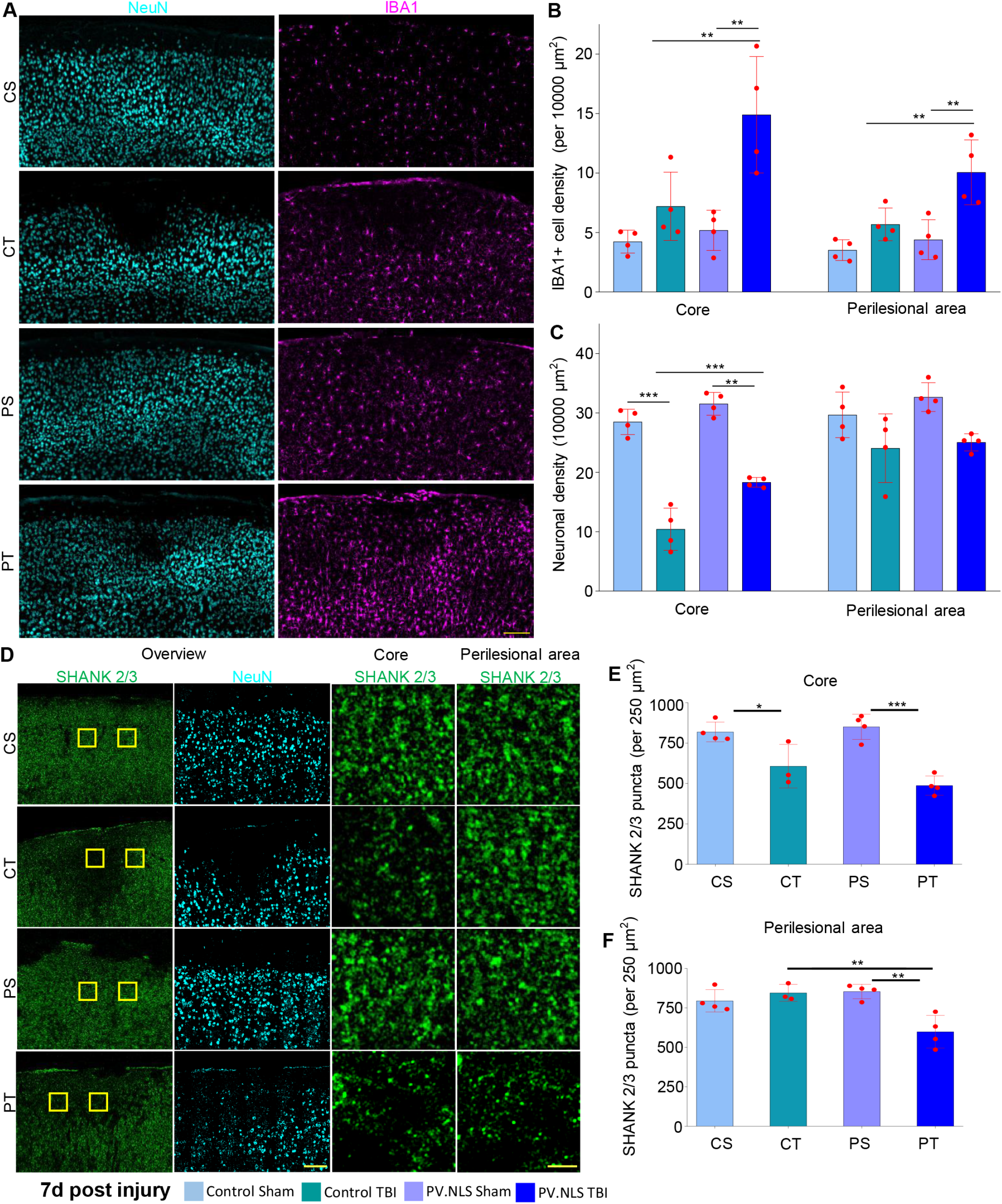
Buffering of neuronal nuclear Calcium enhances subacute microgliosis and synapse loss 7 days post trauma. A) Confocal images of IBA1+ cells from the somatosensory cortex layer 2-3 obtained from Control Sham (CS), Control TBI (CT), PV.NLS Sham (PS) and PV.NLS TBI (PT) mice. Shown are NeuN (cyan) and IBA1 (magenta). Scale bar: 100μm. B) No difference of IBA1+ cells densities in either the core or the perilesional area 7 days post TBI compared to sham (CS vs CT; Core 4.223 ± 0.965 vs 7.200 ± 2.870; Perilesional area 3.517 ± 0.873 vs 5.683 ± 1.374). Buffering of nuclear calcium signaling in TBI significantly increased the density of IBA1+ cells compared to TBI alone (CT vs PT; Core 7.200 ± 2.870 vs 14.900 ± 4.894; Perilesional area 5.683 ± 1.374 vs 10.060 ± 2.728). Data is shown as mean ± SD. N = 4 mice. **: p < 0.01. C) Significant decrease of the density of NeuN+ cells in the core of the injury 7 days post TBI (CS vs CT; 28.500 ± 2.132 vs 10.420 ± 3.555), however not in the perilesional area (CS vs CT; 29.680 ± 3.830 vs 24.060 ± 5.783). Buffering of nuclear calcium signaling in TBI significantly reduced the neuronal loss in the core injury compared to TBI alone (CT vs PT; 10.420 ± 3.555 vs 18.320 ± 0.816), however no difference was observed in the perilesional area (CT vs PT; 24.060 ± 5.783 vs 25.050 ± 1.444). Data are shown as mean ± SD. N = 4 mice. **: p < 0.01; ***: p < 0.001. D) Confocal images from the somatosensory cortex layer 2-3 obtained from Control Sham (CS), Control TBI (CT), PV.NLS Sham (PS) and PV.NLS TBI (PT) mice. Shown are SHANK2/3 (green) and NeuN (cyan). Yellow inserts show analyzed core and perilesional areas. Scale bar 100μm (overview) and 5μm (insert). E) Significant decrease in core synaptic density 24h post TBI compared to sham (CS vs CT; 819.1 ± 61.44 vs 607.1 ± 134.5). Buffering of nuclear calcium signaling did not alter core synaptic density compared to TBI (CT vs PT; 607.1 ± 134.5 vs 423.3 ± 60.1). Data is shown as mean ± SD. N = 4 mice. *: p<0.05; ***: p < 0.001. F) No difference in perilesional synaptic density 24h post TBI compared to sham (CS vs CT; 794.9 ± 70.5 vs 845.2 ± 53.7). Buffering of nuclear calcium signaling significantly decreases synaptic density in the perilesional area compared to TBI (CT vs PT; 845.2 ± 53.7 vs 599.2 ± 103.7). Data is shown as mean ± SD. N = 4 mice. **: p < 0.01

Taken together, these findings demonstrate that enhanced microgliosis resulting from NC buffering following TBI is not transient but is maintained for up to 7 days and is associated with a larger area of synaptic loss, thus linking the robust cellular inflammatory response with synaptic damage.

### 5. Blunting neuronal nuclear calcium worsens acute motor disturbances upon TBI

Next we explored the functional impact of the heightened microgliosis and synaptic loss elicited by NC buffering after TBI. Since the somatosensory area of the brain provides strong excitatory drive to the primary and secondary whisker motor area (Matyas et al., 2010; Yamashita et al., 2018; Commisso et al., 2018), and based on evidence that silencing of the somatosensory area results in decreased whisking (Sreenivasan et al., 2016), we hypothesized that the disruption of synaptic networks in the somatosensory cortex might affect the whisking activity even in absence of a direct lesion of the motor area.

In fact, high-speed recordings of the spontaneous whisking activity of whiskers contralateral to the injury site assessed as number of whisking events (Fig. 6A) revealed that CT mice did show a decrease in spontaneous whisking already at 1 dpi with a further deterioration at 3dpi (nadir point) before showing a trend towards recovery at 7 dpi (Fig. 6B-C, F). Note, mice could not be tested at earlier time points because the stress of brain injury made them characteristically uncooperative. While the whisking activity of PS mice were comparable to CS mice, PT mice displayed a significantly larger decline in whisking at 1 dpi compared to CT mice (Fig. 6C, E-F), indicating a more severe sensorimotor dysfunction in this group. However, the activity of PT mice converged with the CT counterparts by 3 dpi and with a similar recovery at 7 dpi. The kinetic parameters of single whisking events (Supp. Fig. 3A-F) were comparable in the four groups, underscoring the sparing of the motor cortices; likewise, the activity of ipsilateral whiskers, controlled by the contralateral, uninjured side, was also comparable across the four groups (Fig. 6G).

**Figure 6.**
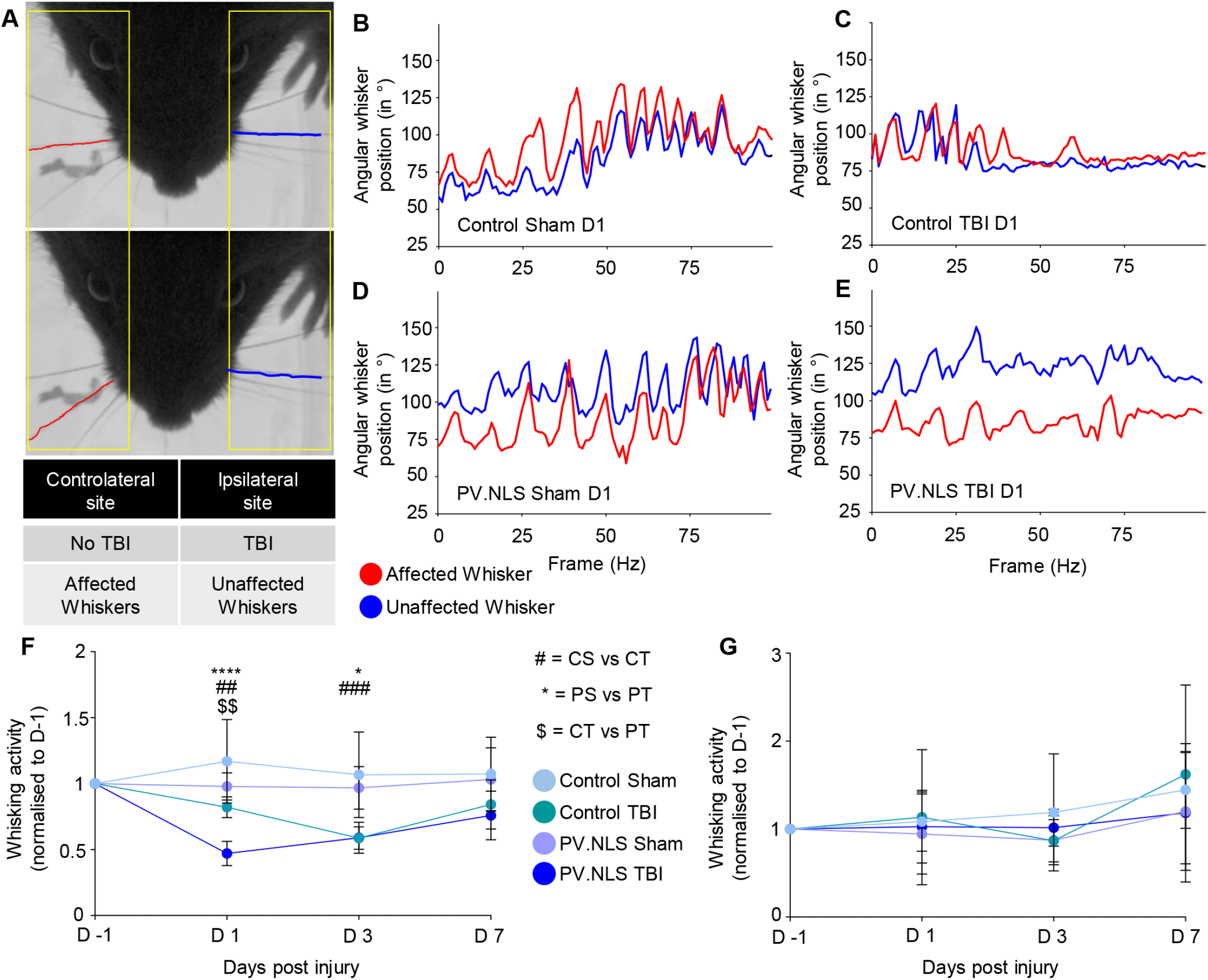
Blunting neuronal nuclear calcium worsens acute motor disturbances upon TBI. A) Representative images of analyzed mouse whiskers, showing the affected (red) and unaffected (blue) whiskers. B)-E) The angular position of the affected whiskers compared to the unaffected whiskers was decreased at D1 post injury in PV.NLS TBI mice, but not in the other treatment groups. F) Significant decrease of whisker activity after nuclear calcium buffering in TBI compared to TBI alone 1d post TBI (CT vs PT; 0.82 ± 0.08 vs 0.47 ± 0.09). 3d after TBI, there was a significant decrease in whisking activity in TBI compared to sham (CS vs CT; 1.17 ± 0.32 vs 0.82 ± 0.08), and buffering of nuclear calcium did not evoke a further decrease at this timepoint (CT vs PT; 0.59 ± 0.09 vs 0.59 ± 0.12). 7d after TBI, the decrease of whisking activity in the Control TBI and PV.NLS TBI groups resolved and moved back towards baseline. N = 8 mice. *: p < 0.05; **: p < 0.01; ****: p < 0.0001. G) No significant changes to the whisking activity of the unaffected whisker was observed. N=8 mice.

Thus, buffering of neuronal NC leads to a more severe acute functional disruption of the injured network.

### 6. Targeted transcriptome analysis reveals new neuronal nuclear calcium-regulated mediators of neuro-glia crosstalk after TBI

In order to gain mechanistic insights into the processes triggering enhanced microgliosis and phagocytic phenotype occurring when neuronal NC is inhibited in TBI, we obtained a targeted nanosting transcriptome analysis of genes involved in neuroinflammatory cascades of the injected/injured area from CS, CT, PS and PT groups at 3h post injury. After pre-processing, quality control and normalization, the Principal Component Analysis (PCA) analysis distinctly defined the four groups (Fig. 7A), indicating significant differences in their transcriptome. Furthermore, the biological replications belonging to the four groups clustered together, segregated by treatment, in the unsupervised hierarchical clustering (not shown). The comparison of the transcriptome of CT vs PT samples (Fig. 7B, C) identified 74 upregulated and 72 downregulated genes (full list in Supp. Table 1). Specifically, PT samples displayed a significant elevation in the expression of gene associated with phagocytic activity (*CD68, Lamp1, Lamp2, Itgam, Ctss, Tlr2, Ifi30*), antibody-mediated phagocytosis (*Fcgr1, Fcgr2b, Fcgr3, Fcer1g*) and in particular with DAM (*TREM2, Clec7a, Cst7, complement C4a, ApoE, Cx3cr1, Csf1r, Spp1, Tyrobp, Grn*). Furthermore, it showed a distinct elevation in the transcription of several other complement factors of the classical pathway (*C1qa, C1qb, C1qc, C3*), interferon-response genes (*Irf7, Irf8, STAT1*), chemokines and other migration factors (*Ccl3, Ccl5, Cxcl9, Cxcl10*). Thus, PT samples displayed a transcriptome compatible with the histological evidence of increased microglial recruitment as early as 3h after injury, with a DAM and phagocytic phenotype.

**Figure 7.**
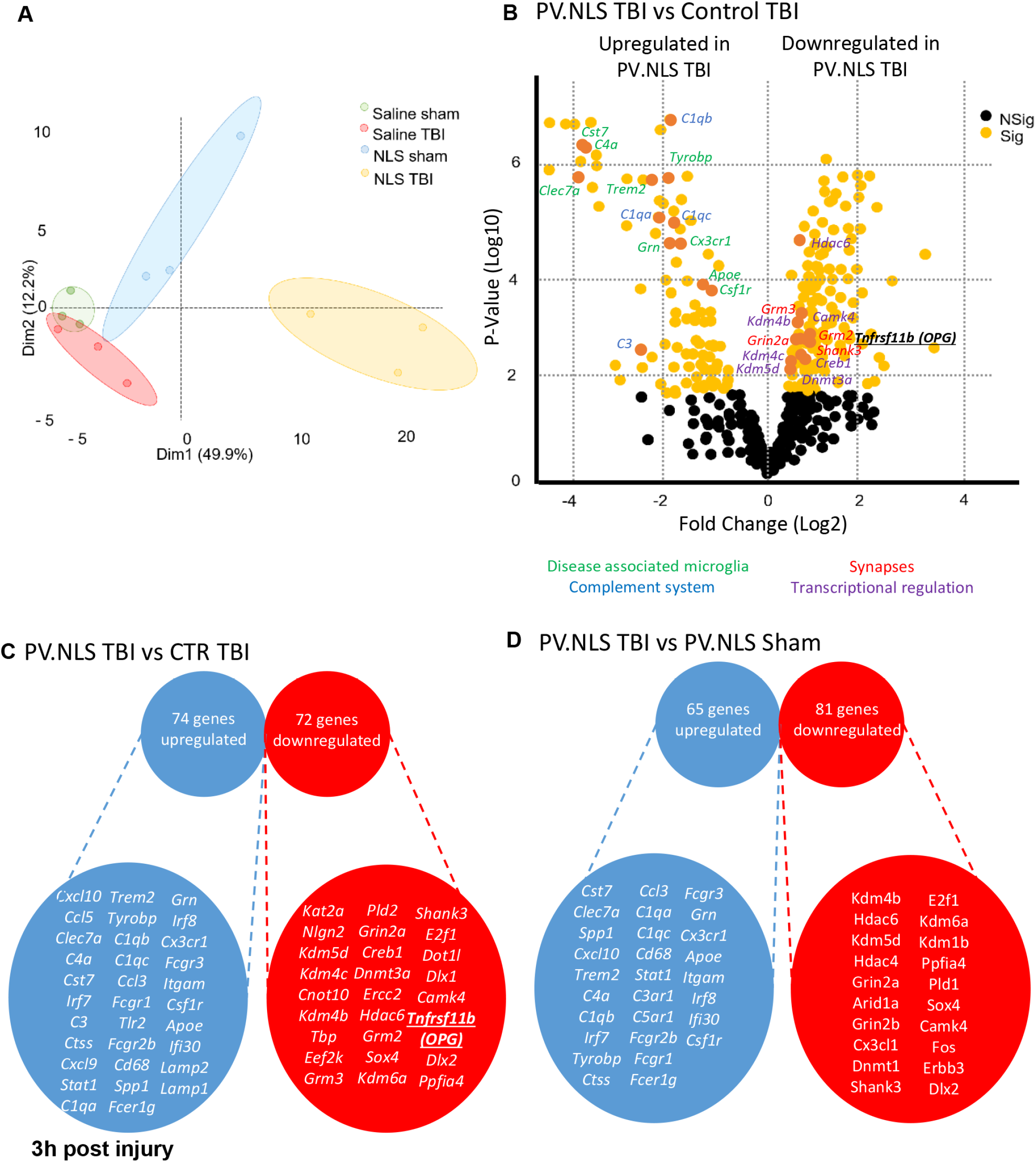
Targeted transcriptome analysis reveals new neuronal nuclear calcium-regulated mediators of neuro-glia crosstalk after TBI. A) Post preprocessing principal component analysis (PCA) plot showing distinct clustering of each group with Dim1 (49.9%) and Dim2 (12.2%) confidence ellipses around the groups. PV.NLS TBI shows no overlap with other groups; other groups show minimal overlap. B) Volcano Plot of PV.NLS TBI vs control TBI shows distinct upregulation of genes related to disease associated microglia (green) or complement system (blue) and a distinct down regulation of genes related to synaptic function (red) and transcriptional regulation (purple). C) Transcriptional analysis shows 74 genes upregulated and 72 genes downregulated in the PV.NLS TBI vs control TBI groups. A subset of genes is depicted. D) Transcriptional analysis shows 65 genes upregulated and 81 genes downregulated in the PV.NLS TBI vs PV.NLS sham group. A subset of genes is depicted. A complete list of significantly differentially expressed genes can be found in supplementary information.

On the other hand, a number of genes associated with synaptic proteins (*Shank3, Grin2a, Grm2, Grm3*) or involved in transcriptional and epigenetic regulation (*Kdm4b and c, Kdm5d, Dnmt3a, Hdac6, CamKIV and Creb1*) were downregulated. Very few soluble mediators were identified among the downregulated genes. One of these, *TNFRSF11b,* also known as Osteoprotegerin (OPG), encodes a soluble mediator involved in suppressing the function of phagocytes, in bone and connective tissue (Glasnovic et al., 2020), and was then selected for further investigation.

The comparison of the PT and PS transcriptomes revealed 65 upregulated and 81 downregulated genes (Fig. 7D). The list of upregulated and downregulated genes was remarkably similar to those that were revealed by the comparison of PT and CT transcriptomes, with a selective upregulation of complement-associated and DAM-associated genes (among others, *Clec7a, Cst7, C4a, Spp1, ApoE* as well as *C5ar1, C1qb, C1qa, C3ar1*), antibody-dependent phagocytosis and interferon-response genes. Among the downregulated genes, synaptic proteins and epigenetic regulators were also prominently represented. Taken together these findings suggest that the induction of DAM-like transcriptional signatures and downregulation of synaptic genes is not a consequence of NC blockade alone but a fingerprint of TBI occurring in the context of NC blockade.

### 7. Neuronal expression of osteoprotegerin is upregulated by nuclear calcium signaling and neuronal activity in TBI

OPG is a decoy receptor for RANKL (Glasnovic et al., 2020) and has been previously implicated in the regulation of osteoclasts. In the bone of OPG^-/-^ mice osteoclasts are overactive and OPG overexpression suppresses their phagocytic activity (Chagraoui et al., 2003). Interestingly, OPG also restricts microglial reactivity to bacterial infection (Kichev et al., 2017).

In this contect, we used single molecule *in situ* mRNA hybridization to confirm the modulation of OPG expression and pinpoint the cellular source of OPG. OPG mRNA colocalized with NeuN and VGLUT2 in all four treatment groups, indicating a neuronal source (Fig. 8A). Notably, the number of OPG mRNA molecules increased in neurons 3h after TBI (Fig. 8A, C-D) but this upregulation was abolished in neurons expressing PV.NLS.mCherry (Fig. 8A, C-D).

**Figure 8.**
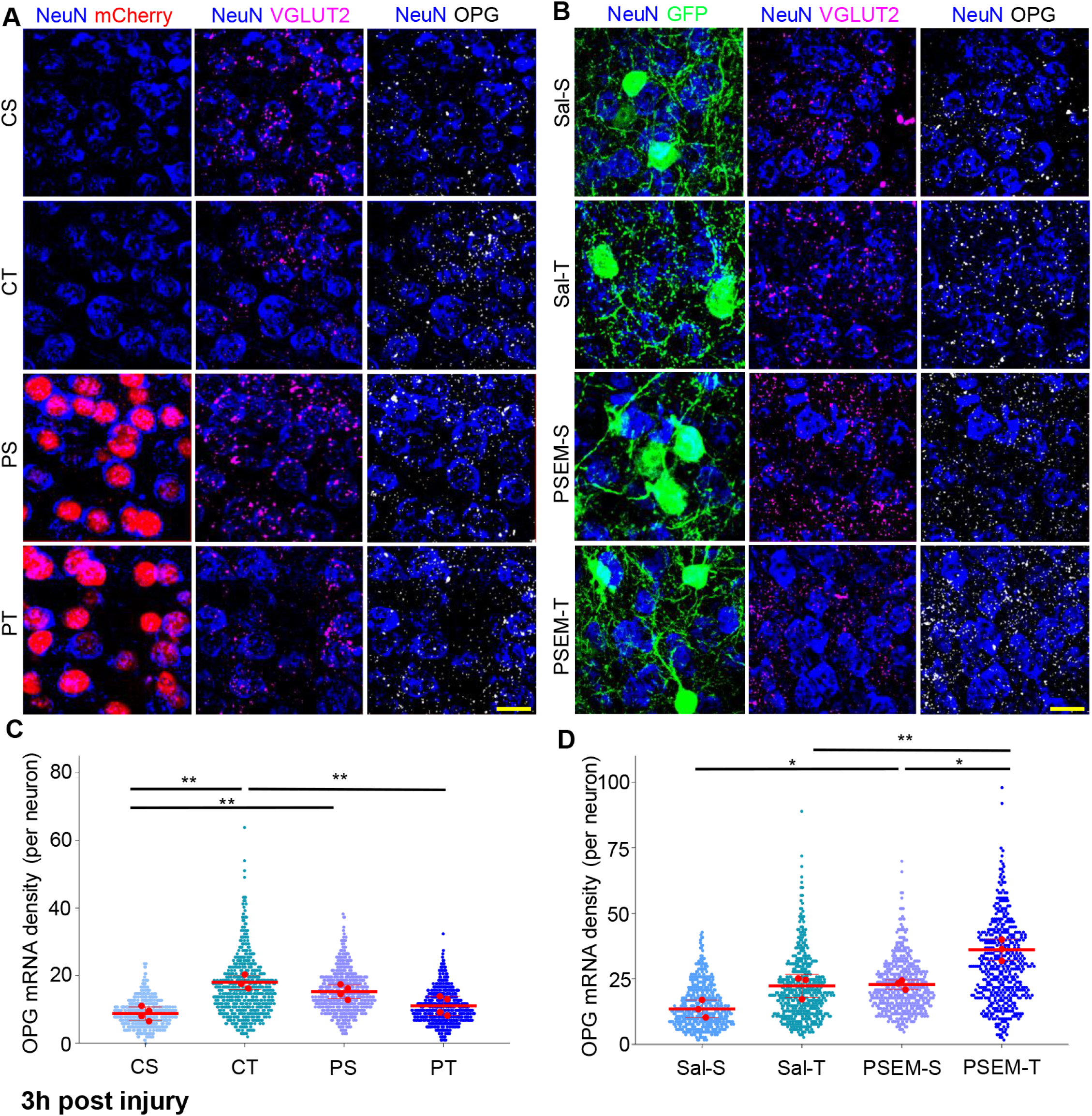
Neuronal expression of osteoprotegerin is upregulated by nuclear calcium signaling and neuronal activity in TBI. A) Confocal images from the somatosensory cortex layer 2-3 obtained from Control Sham (CS), Control TBI (CT), PV.NLS sham (PS) and PV.NLS TBI (PT) mice. In situ hybridisation was performed for OPG (TNFRSF11b; white) and VGLUT2 (magenta), co-staining was performed with NeuN (blue) and mCherry (red). Scalebar: 2Oμm. B) Confocal images from the somatosensory cortex layer 2-3 obtained from Saline Sham (Sal-S), Saline TBI (Sal-T), PSEM sham (PSEM-S) and PSEM TBI (PSEM-T) mice. In situ hybridisation was performed for OPG (TNFRSF11b; white) and VGLUT2 (magenta), co-staining was performed with NeuN (blue) and GFP (green). Scalebar: 2Oμm. C) Significant increase of OPG mRNA density 3h post TBI compared to sham (CS vs CT; 8.786 ± 1.958 vs 18.03 ± 2.085). Buffering of nuclear calcium in TBI significantly decreases OPG mRNA density compared to TBI alone (CT vs PT; 18.03 ± 2.09 vs 11.06 ± 2.80). Data is shown as a scatterplot with the mean of each animal indicated in red with mean ± SD. The mean of each animal was considered for statistical calculations. N=4 mice. **: p < 0.01. D) Significant increase of OPG mRNA density 3h post TBI compared to sham (Sal-S vs Sal-T; 13.46 ± 3.339 vs 22.24 ± 4.432). Chemogenetic inhibition of PV interneurons in TBI significantly increases OPG mRNA density compared to TBI alone (Sal-T vs PSEM-T; 22.24 ± 4.43 vs 31.77 ± 4.10). Data is shown as a scatterplot with the mean of each animal indicated in red with mean ± SD. The mean of each animal was considered for statistical calculations. N = 3 mice. *: p < 0.05; **: p < 0.01.

We then wondered whether OPG expression could be enhanced by increasing neuronal firing through chemogenetic approaches. To achieve this goal, we injected an AAV encoding the inhibitory PSAM(Gly)-GFP construct into PV-Cre mice. PSAM(Gly)-GFP was successfully expressed in PV+ interneurons (Fig. 8B). Upon PSEM administration, PSAM(Gly) decreased the excitability of PV interneurons, thereby reducing perisomatic inhibition of neighboring excitatory neurons and resulting in their increased firing (Chandrasekar et al., 2019). Reduced PV firing (PSAM(Gly)+PSEM) resulted in the upregulation of OPG in sham mice and even more robustly in TBI mice, compared to control mice (PSAM(Gly)+vehicle; Fig. 8B, E-F)).

So it can be concluded that OPG is upregulated in neurons upon TBI through a process involving neuronal firing and NC signals.

### 8. Re-expression of OPG in neurons with nuclear calcium buffering reduces microgliosis and prevents synaptic degradation after TBI

To establish a mechanistic link between neuronal OPG and TBI-induced microgliosis, we reexpressed OPG in neurons by NC blockade. The AAV expressing OPG under the hSyn promoter was sufficient to produce a massive upregulation of OPG expression irrespective of PV.NLS expression (Supp. Fig. 4A-B). Mice were injected with a mix of AAV encoding either PV.NLS alone (PT) or PV.NLS and OPG (POT) 30 days before being subject to TBI. Two more groups were considered, both injected with AAV encoding an empty control vector, and subject to sham surgery (CS) or TBI (CT). Re-expression of OPG substantially decreased the extent of microgliosis as determined by the density of IBA1+ and IBA1+/TMEM119+ cells (Fig. 9A-B) and IBA1+/CD11c+ cells after TBI in POT mice compared to PT mice (Fig. 9A-B). Furthermore, OPG re-expression substantially decreased the proportion of cells expressing CST7 (Fig. 9A, C) suggesting that OPG re-expression is responsible for the decrease in microglial recruitment and for the reduced expression of this DAM marker.

**Figure 9.**
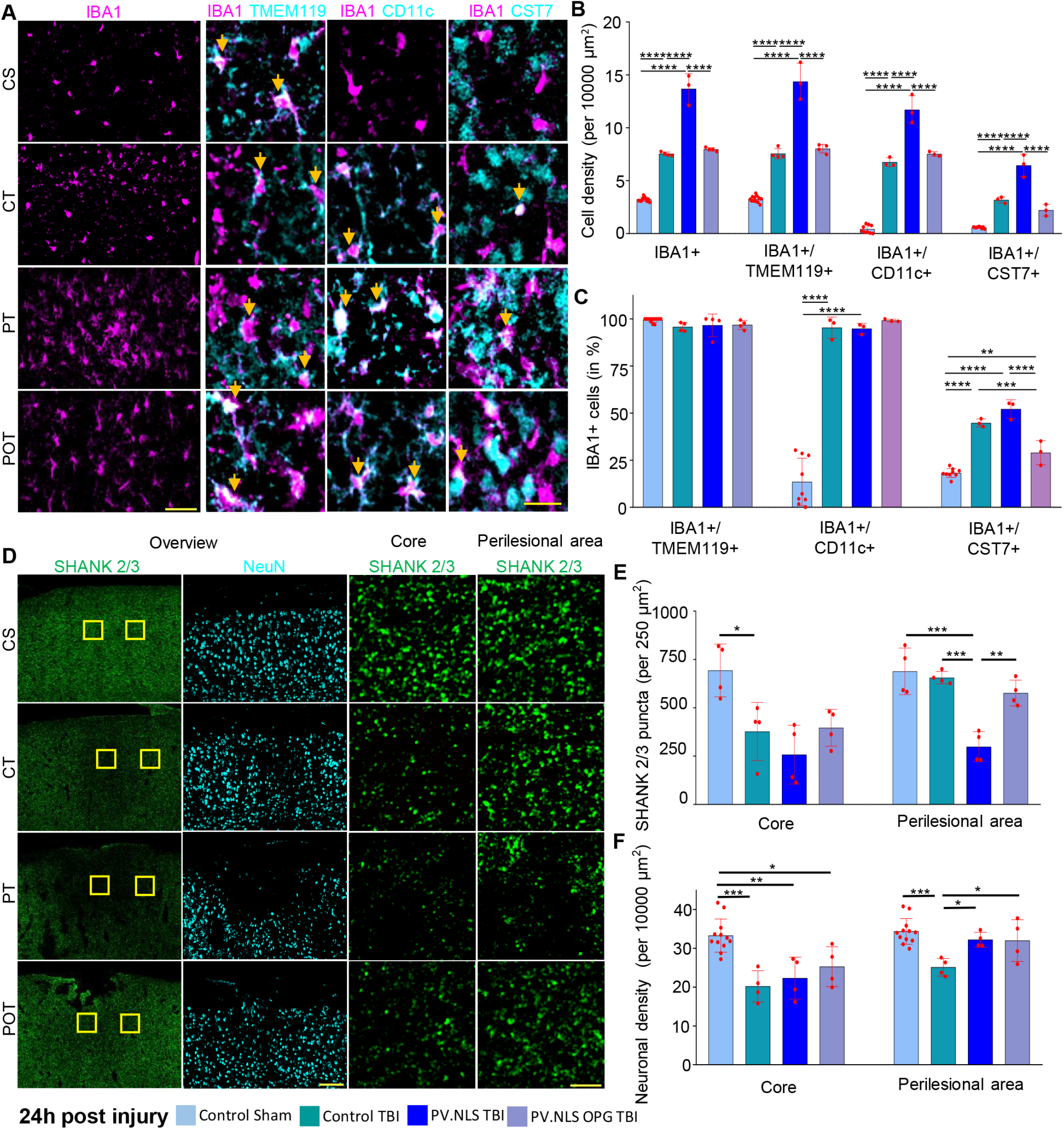
Re-expression of OPG in neurons with nuclear calcium buffering reduces microgliosis and prevents synaptic degradation after TBI. A) Confocal images of somatosensory cortex layer 2-3 obtained from Control Sham (CS), Control TBI (CT), PV.NLS TBI (PT) and PV.NLS OPG TBI (POT) mice. Shown are IBA1 (magenta), TMEM119 (cyan), CD11c (cyan) and CST7 (cyan). Scale bar: 40μm (overview) and 20μm (insert). B) Significant increase of IBA1+/TMEM119+, IBA1+/CD11c+ and IBA1+/CST7+ cells 24h post TBI compared to sham (CS vs CT; for IBA1+/TMEM119+ 3.251 ± 0.297 vs 7.560 ± 0.486; for IBA1+/ CD11c+ 0.418 ± 0.376 vs 6.746 ± 0.427; for IBA1+/CST7+ 0.577 ± 0.077 vs 3.175 ± 0.291). Buffering of nuclear calcium signaling significantly increased the density of IBA1+/TMEM119+, IBA1+/CD11c+ and IBA1+/CST7+ cells 24h after TBI compared to TBI alone (CT vs PT; for IBA1+/TMEM119+ 7.560 ± 0.486 vs 14.370 ± 1.745; for IBA1+/CD11c+ 6.746 ± 0.427 vs 11.700 ± 1.330; for IBA1+/CST7+ 3.175 ± 0.291 vs 6.440 ± 1.053). Re-expression of OPG together with buffering of nuclear calcium signaling significantly decreased IBA1+/TMEM119+, IBA1+/CD11c+ and IBA1+/CST7+ cells compared to PV.NLS TBI (PT vs POT; for IBA1+/TMEM119+ cells 14.37 ± 1.745 vs 8.024 ± 0.409; for IBA1+/CD11c+ cells 11.7 ± 1.33 vs 7.524 ± 0.2182; for IBA1+/CST7+ cells 6.444 ± 1.053 vs 2.206 ± 0.548). Data is shown as mean ± SD. N = 4 mice. ***: p<0.001; ****: p < 0.0001. C) No difference in percentage of IBA1+ cells also positive for TMEM119+ between treatment groups (CS = 99.400 ± 1.154%; CT = 95.750 ± 2.341%; PT = 96.610 ± 5.989%; POT = 96.830 ± 2.378%). Significant increase in percentage IBA1+ cells also positive for CD11c+ and CST7+ 24h post TBI compared to sham (CS vs CT; for CD11c+ 13.490 ± 12.530% vs 95.360 ± 5.657%; for CST7+ 18.240 ± 2.379% vs 44.790 ± 2.122%). Buffering of nuclear calcium signaling in TBI did not alter the percentage of IBA1+ cells copositive for CD11c+ and CST7+ compared to TBI alone (CT vs PT; for CD11c+ 95.360 ± 5.657% vs 94.850 ± 2.757%; for CST7+ 44.790 ± 2.122% vs 52.140 ± 4.91%). Reexpression of OPG together with buffering of nuclear calcium signaling did not alter the percentage of IBA1+ cells copositive for CD11c+ compared to PV.NLS TBI (PT vs POT; 94.850 ± 2.757% vs 99.170 ± 0.721%), but decreased the percentage of IBA1+ cells copositive for CST7+ compared to PV.NLS TBI (PT vs POT; 52.140 ± 4.910% vs 28.980 ± 6.444%). Data is shown as mean ± SD. N = 3 mice. **: p < 0.01; ***: p < 0.001; ****: p < 0.0001. D) Confocal images of somatosensory cortex layer 2-3 obtained from Control Sham (CS), Control TBI (CT), PV.NLS TBI (PT) and PV.NLS OPG TBI (POT) mice. Shown are SHANK2/4 (green) and NeuN (cyan). Yellow inserts show analyzed core and perilesional areas. Scale bar: 100μm (overview) and 5μm (insert). E) Significant decrease in core synaptic density 24h post TBI compared to sham (CS vs CT; 693.1 ± 136.8 vs 377.2 ± 149.9). Buffering of nuclear calcium signaling did not alter core synaptic density compared to TBI (CT vs PT; 377.2 ± 149.9 vs 257.4 ± 152.0). Re-expression of OPG together with buffering of nuclear calcium signaling did not alter core synaptic density compared to PV.NLS-TBI (PT vs POT; 257.4 ± 152.0 vs 393.3 ± 95.3). No difference in perilesional synaptic density 24h post TBI compared to sham (CS vs CT; 688.4 ± 120.5 vs 655.5 ± 32.0). Buffering of nuclear calcium signaling significantly decreases synaptic density in the perilesional area compared to TBI (CT vs PT; 655.5 ± 32.0 vs 298.3 ± 78.1). Re-expression of OPG together with buffering of nuclear calcium signaling significantly increased the synaptic density in the perilesional area compared to PV.NLS-TBI (PT vs POT; 298.3 ± 78.1 vs 576.3 ± 66.8). Data is shown as mean ± SD. N = 4 mice. *: p < 0.05; **: p < 0.01; ***: p < 0.001. F) Significant decrease in core neuronal density 24h post injury compared to sham (CS vs CT; 33.27 ± 4.27 vs 20.21 ± 4.03). Buffering of nuclear calcium signaling or OPG re-expression did not alter core neuronal density compared to TBI alone (CT vs PT; 20.21 ± 4.03 vs 22.32 ± 5.40; CT vs POT; 20.21 ± 4.03 vs 25.28 ± 5.14). TBI alone significantly reduces neuronal density in the perilesional area compared to sham, nuclear calcium buffering and OPG re-expression (CS vs CT 34.35 ± 3.32 vs 25.13 ± 2.26; CT vs PT 25.13 ± 2.26 vs 32.2 ± 1.91; CT vs POT 25.13 ± 2.26 vs 32.01 ± 5.38). Data is shown as mean ± SD. N=4 mice. *: p < 0.05; **: p < 0.01; ***: p < 0.001.

Re-expression of OPG also normalized the synaptic density in the perilesional area, resulting in a smaller area of synaptic loss (Fig. 9D, E). It is worth noting that synaptic loss in the core was not affected by OPG expression, indicating that the damage caused by TBI was comparable in all injured groups. While neuronal density in the core of POT samples was similar in CT and PT samples, further confirming the reproducibility of TBI, and was comparable to PT in the perilesional area (Fig. 9D, F), indicating that while microgliosis and synaptic loss may be influenced by OPG, other aspects of the TBI pathophysiology may not be restored once the mechanical damage has occurred.

Thus, re-expression of OPG in neurons reverted the increased microgliosis and more extensive synaptic loss observed upon trauma when NC signaling was inhibited.

### 9. OPG concentration increases in CSF samples from human TBI patients

Finally, we explored the translational value of OPG role in human TBI patients. We considered series of CSF samples obtained from patients with severe TBI on the day of trauma (D0) and 1, 3 and 7 days later (D1, D3 and D7, respectively). Unfortunately, complete longitudinal CSF samples were not available and therefore distinct cohorts were considered. Samples from 6 non-TBI patients who required neurosurgery were included as baseline controls (clinical and demographic data are detailed in Table 1). The clinical cohort from which these samples were obtained has been the object of previous work (Yan et al., 2014). The four TBI cohorts (D0, D1, D3, D7) did not differ in terms of age, Glasgow Coma Scale (GCS) at admission, Injury Severity Score (ISS) at admission and Glasgow Outcome Scale-Extended (GOSE) at 6 months (Table 1); the age of the individuals in the control group was significantly different from those of the cases.

**Table 1.** Clinical and demographic characterization of human TBI patient (Melbourne Cohort).

|  | **Control group\*** | **D0 group** | **D1 group** | **D3 group** | **D7 group** | **p value** |
| --- | --- | --- | --- | --- | --- | --- |
| N (M/F) | 6 (2/4) | 18 (12/6) | 18 (15/3) | 15 (11/4) | 16 (12/4) |  |
| Age (median, range) | 71 (53-85) | 35 (23-54) | 36 (23-48) | 31 (19-66) | 28.5 (19-66) | 0.43 (K-W) |
| Glasgow Coma Scale (median, range) | N/A | 5.5 (3-10) | 5.5 (3-10) | 6.5 (3-10) | 5.5 (3-11) | 0.88 (K-W) |
| Injury Severity Score (median, range) | N/A | 39 (17-50) | 35 (18-59) | 33.5 (17-58) | 34.5 (17-50) | 0.98 (K-W) |
| Glasgow Outcome Scale-Extended (median, range) | N/A | 3 (1-8) | 4 (1-8) | 4 (1-8) | 3.5 (1-8) | 0.97 (K-W) |

\* non-TBI CSF samples were obtained from hydrocephalus patients (without previous TBI) undergoing implantation of ventriculoperitoneal shunts.

Compared to controls, levels of OPG at D0 were significantly elevated and displayed further increase at D1. Which returned to baseline at D3 and D7 (Fig. 10).

**Figure 10.**
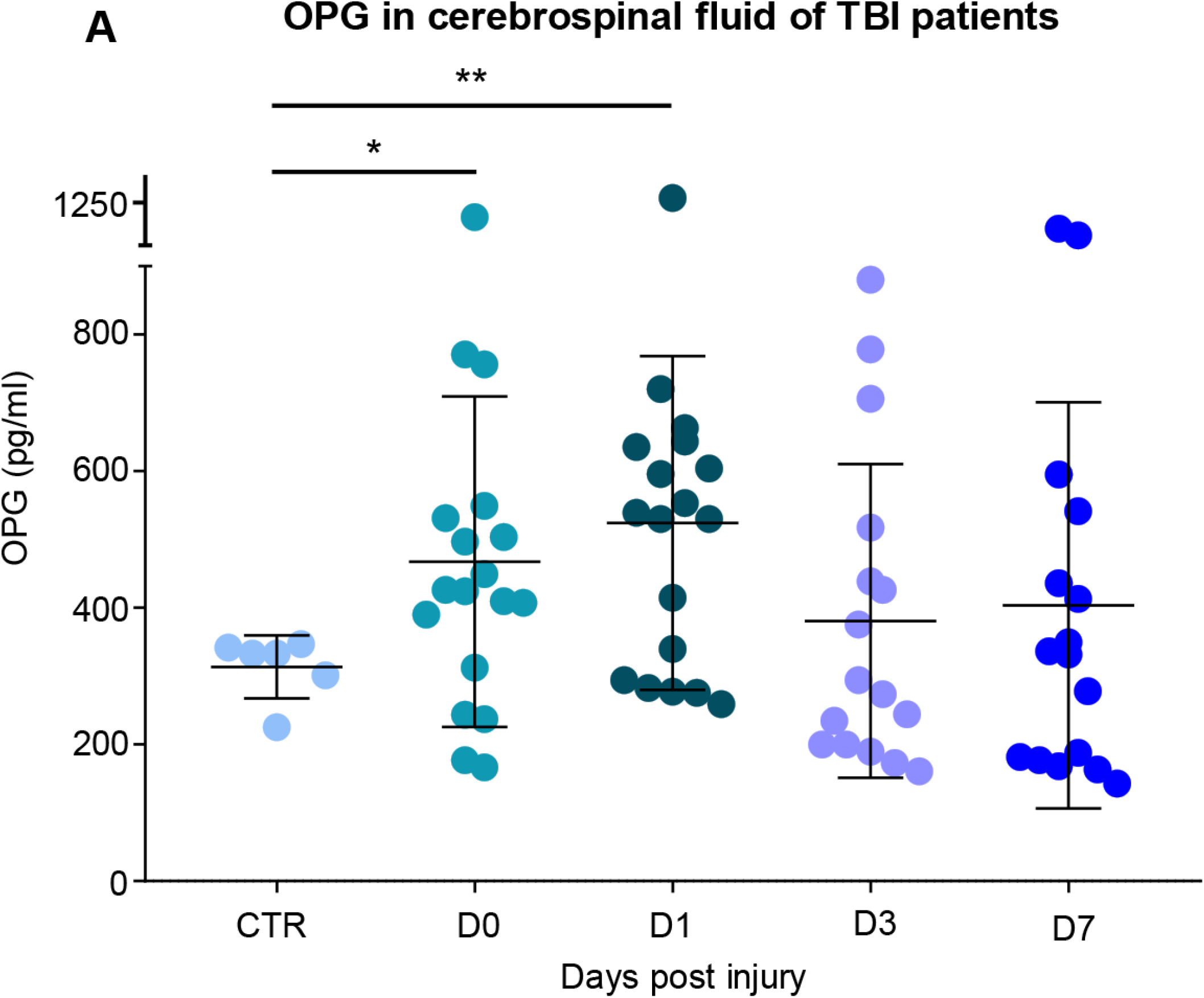
OPG concentration increases in CSF samples from human TBI patients. A) Significant increase in osteoprotegerin (OPG) levels in cerebrospinal fluid (CSF) of TBI patients upon admission (CTR vs D0; 313.0 ± 46.1 pg/ml vs 467.5 ± 241.0 pg/ml) and 1d after trauma (CTR vs D1; 313.0 ± 46.1 pg/ml vs 524.1 ± 244.3 pg/ml). Expression levels decreased subsequently 3d (CTR vs D3; 313.4 ± 46.1 pg/ml vs 380.7 ± 229.6 pg/ml) and 7d (CTR vs D7; 313.4 pg/ml vs 403.6 ± 297.3 pg/ml) after trauma. Data is shown as mean ± SD. CTR N = 6; D0 N = 18; D1 N = 18; D3 N = 16; D7 N = 16. *: p < 0.05; **: p < 0.01.

This data supports a robust elevation of OPG in human CSF, with a peak in the early phase and followed by a rapid normalization, suggesting that OPG upregulation may have a pathophysiological role not only in rodent but also in human TBI.

## Discussion

In this study we have demonstrated that neuronal NC signals are, unexpectedly, critical in keeping in check the activation of microglia following a mild TBI. Buffering of NC does not result in increased neuronal loss but rather a much greater recruitment of microglia displaying a typical disease-associated phenotype. Heightened microglia activation and accumulation was concomitant to a more profound loss of synapses, ultimately linked to a worse functional impairment. For the first time, we identify OPG as a neuronally-originated, activity-dependent mediator of neuron-microglial crosstalk and an important player in restricting microglial reactivity and synapse elimination after TBI (Fig. 11). The translational relevance of our findings is underscored by the elevation in OPG levels in CSF of TBI patients occurring within the first 24 h after trauma.

**Figure 11.**
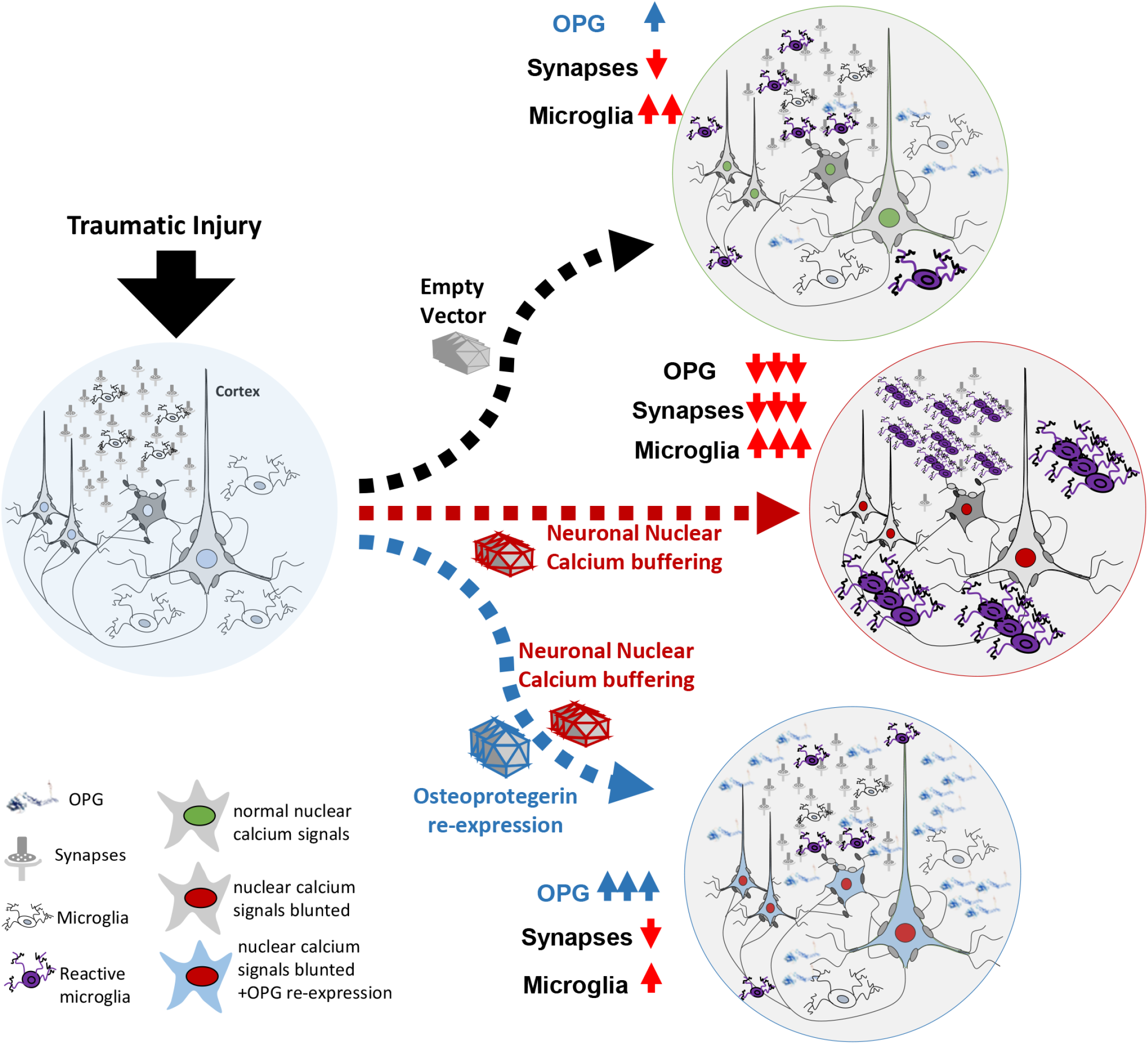
Neuronal nuclear calcium signaling controls neuroinflammation in TBI. In normal conditions (empty vector), neuronal activity induces a transcriptional program that limits the induction of reactive microglia upon TBI and the associated loss of synapses. When neuronal nuclear calcium is inhibited, however, synaptic loss and microglial recruitment are strongly increased upon TBI, but these effects are reversed by the re-expression of Osteoprotegerin (OPG).

A number of soluble and membrane-bound molecules have been reported to mediate the interaction between neurons and microglia (Szepesi et al., 2018). Neuronal CD200 is involved in maintaining microglia in a non-reactive state by engaging the CD200L expressed on microglia (Hoek et al., 2000). Similarly, in spinal cord injury the suppression of CD200/CD200L worsened the functional outcome (Lago et al., 2018). Neuronal membrane-bound fractalkine and CD47 have also been implicated in restricting microglial reactivity and preventing the phagocytosis of neurons (Biber et al., 2007). Several soluble mediators are released by neurons resulting in an impact on microglia function, particularly maintaining a non-reactive state. Among these, BDNF appears to exert a suppressive signal on microglia, since the loss of BDNF results in microgliosis (Onodera et al., 2021). On the other hand, complement factors secreted by neurons have been shown to enhance microglial reactivity and induce microglia-mediated elimination of synapses (Lim et al., 2021; Hammond et al., 2020; Simonetti et al., 2013; Vasek et al., 2016). These factors are either dependent on neuronal activity (such as BDNF) or appear to be dysregulated as shown in our transcriptome study (complement factors), suggesting that NC is a central hub in determining not only the overall survival of neurons, but also the mobilization of local microglia upon injury.

The unbiased screening of transcripts altered by NC buffering in the injured brains revealed a number of proteins involved in inducing a reactive microglia state. Among the few soluble mediators detected in the screening, we selected OPG for further study. OPG is a member of the TNF superfamily and it is more specifically part of the RANKL/RANK/OPG axis (Glasnovic et al., 2020; Hanada, 2021). RANK is the cognate receptor of the RANK ligand (RANKL), whereas OPG is a soluble decoy that binds to RANKL, effectively, downregulating RANK signaling. OPG was first identified for its function in bone metabolism: the interaction of RANK with RANKL is a critical inducer of multinucleated osteoclasts involved in bone resorption. In fact, OPG prevents the RANK signaling and reduces osteoclast formation (Walsh and Choi, 2014). OPG expression in the brain has been reported previously (Hofbauer et al., 2004; Kichev et al., 2017), but its role in microglial biology is currently controversial. OPG has been shown to be upregulated by ischemia in juvenile rats and mice (Kichev et al., 2014; Shimamura et al., 2014) and is elevated in the serum of stroke patients (Jensen et al., 2010). Most notably, in a murine model of middle-cerebral artery occlusion, OPG deletion results in decreased infarct volume and reduced brain oedema (Shimamura et al., 2014). Furthermore, OPG^-/-^ mice were reported to display lower levels of inflammatory cytokines, such as IL-6 and TNF-α, and diminished infiltration of macrophages around the infarct area. In this stroke study, it was concluded that OPG signaling has an overall detrimental effect, such as decreasing the RANK/RANKL signaling in macrophages and neurons and leading to a pro-inflammatory phenotype in macrophages and consequent increased neuronal vulnerability (Shimamura et al., 2014). On the other hand, OPG levels are reported to be reduced in multiple sclerosis (MS), a condition characterized by CNS inflammation (Glasnovic et al., 2018). So, it is speculated that OPG signaling may be protective in this autoimmune disease, by decreasing the activity of the RANK/RANKL (Guerrini et al., 2015). Although the role of OPG in neuroinflammation deserves further investigation, which may be highly context-dependent, it is worth noting that in non-cerebral tissues OPG has been shown to reduce inflammation. In fact, dendritic cells from OPG^-/-^ mice produce higher levels of inflammatory cytokines *in vitro* and *in vivo* upon LPS treatment (Chino et al., 2009) and are more effective in driving T-cell responses (Yun et al., 2001). Conversely, OPG administration ameliorates intestinal inflammation and mucosal dendritic cells infiltration (Ashcroft et al., 2003), while OPG overexpression reduces dendritic cell activation in an asthma model (Yang et al., 2019). Our data show that OPG downregulation is at least one of the factors leading to increased microglial reactivity when neuronal NC signaling is inhibited, since re-establishing OPG expression normalized microglial response to mild TBI and prevented the excessive loss of synapses. These findings are strongly consistent with the broader interpretation of OPG as a down regulator of immune activation (Walsh and Choi, 2021).

The relationship of neuronal function with NC signaling as well as local microglial reactivity supports the concept of sustaining healthy neurons and neuronal NC signaling through a targeted, early rehabilitative intervention, which may contribute to dampening the neuroinflammatory response elicited by TBI. In fact, the loss of activity-dependent transcription factor ATF3 results in increased inflammatory infiltrates (Förstner et al., 2018). Moreover, exercise has been shown to have a beneficial effect on TBI-associated microglial activation but, surprisingly, only when delayed; acute exercise appears to increase cytokine levels and microglial reactivity (Piao et al., 2013). Conversely, early exercise does decrease neuroinflammation after intracerebral hemorrhage (Tamakoshi et al., 2020). Since cortical neurons are hypoexcitable in the acute phase of TBI, but hyperexcitable later on (Carron et al., 2016), exercise may not be as effective in inducing neuronal activity-dependent outcomes. Thus, new approaches to early rehabilitation and sensory stimulation are needed to evoke neuronal NC signaling and mitigate local neuroinflammatory response to TBI.

The observed recruitment of reactive microglia following the loss of neuronal NC may have implications beyond the pathophysiology of TBI. In fact, synapse elimination and loss of excitatory inputs are the early and most prominent responses to viral infections, occurring through T-cell induced STAT1 phosphorylation in neurons (di Liberto et al., 2018) and through complement-mediated opsonization and clearance mechanisms (Vasek et al., 2016). Upon synapse elimination, loss of neuronal NC signaling may further induce the local neuroinflammatory response for the removal of the infectious agent. A similar set of mechanisms may be involved in microglial activation occurring in chronic neurodegenerative conditions. In fact, loss of physiological neuronal firing appears to be a shared phenomenon to several degenerative conditions (Roselli and Caroni, 2015), in which microglial cells assume a DAM phenotype (Keren-Shaul et al., 2017; Krasemann et al., 2017). Thus, the role of neuronal NC in maintaining the homeostasis of local inflammatory cells may have a broader regulatory role. Moreover, therapeutic interventions aimed at sustaining a physiological neuronal activity after trauma may be critical in protecting the integrity of synaptic networks by preventing excessive microglial reactivity.

In conclusion, our data show that neurons displaying a normal NC signaling and sustained levels of neuronal function secrete OPG and other factors that restrain the local activation of microglia and the appearance of the DAM-like microglial phenotype. The overall physiological goal may therefore be to single out healthy neurons from the damaged ones and prevent the phagocytosis of the functioning neurons while enhancing it for those that have been damaged by the injury. This mechanism may have broader implications for inflammatory and neurodegenerative diseases characterized by disturbances in neuronal activity and NC signaling.

## Materials and Methods

### Animals

B6;129P2-Pvalb^tml(cre)Arbr^/J mouse line (Jax stock #008069) was a courtesy of Pico Caroni and Silvia Arber. All experiments were approved by the Tierforschungszentrum Ulm and by the Regierungspraesidium Tubingen with licence no. 1420. Only male mice were used to minimize the confounding effect of hormonal oscillations.

The PV-Cre animals were housed in either Type II long IVC cages or open cage in a group of 2-5 with ad libitum access to food and water, nest-building material and a day-night cycle of 12h from 6:00 to 18:00.

### Virus Production

AAV2/9 were prepared as previously described using the iodixanol density gradient protocol (Commisso et al. 2018). The helper plasmids pAd-DeltaF6 (James M. Wilson, Addgene 112867) and p5E18V2/9 (a kind gift from J. Kleinschmidt) were used for the production. The viral suspension was concentrated to a final volume of 200μl and the titre (number of viral genomes/ml) was confirmed by qPCR. The following constructs were expressed: an empty vector hSyn.NLS.myc (negative control); hSyn-PV.NLS.mCherry, (previously reported by Pusl et al. 2002); hSyn-CaMBP4.flag.mCherry (described by Schlumm et al. 2013); hSyn-TNFRSF11b-P2A-mCherry (Vigene biosciences); pAAV(9)-pCAG-A7-floxed-PSAM(L141F, Y115F)-GlyR-GFP-WPRE (inhibitory PSAM(GLY)-GFP construct; Vector Biolabs). The full list of constructs can be found in Supp. Table 2.

### Intracerebral Injection of AAV

Intracerebral injection of the AAV suspension was performed in mice aged p30-35 in correspondence of the primary somatosensory cortex, as previously described (Chandrasekar et al., 2019). Briefly, mice were administered 0,05mg/kg Buprenorphin and 1,0mg/kg Meloxicam 30 minutes before being subject to sevoflurane anesthesia (5% in 95% O_2_) and then positioned in the stereotaxic frame. After the incision of the scalp, a burr hole was drilled in the parietal bone. The AAV suspension (1μl, diluted 1:1 with 1% Fast-Green in DMEM) was injected at the coordinates x=+2.0, y=-2.0, z=-0.2/0.6 using a pulled-glass capillary connected to a Picospritzer microfluidic apparatus. The injection procedure lasted approx. 10 minutes; the capillary was left in place for another 10 minutes. The scalp was then sutured with Prolene 7.0. Mice were administered with a daily dose of 1 mg/kg meloxicam and three daily doses of 0,05 mg/kg Buprenorphin for the next 3 days.

### Traumatic brain injury procedure

Traumatic Brain Injury (TBI) was performed as previously reported (Chandrasekar et al., 2019; olde Heuvel et al., 2019; Flierl et al., 2009). Animals were subjected to TBI at the age of P60-70. Briefly, animals were given sevoflurane anesthesia (4% Servofluran, 96% O_2_) and a dose of 0,05mg/kg buprenorphine. The scalp was incised on the midline and the parietal bone exposed; the animals were then positioned in the TBI apparatus, while under anesthesia, so that the impact site corresponded to the injection site. The impact parameters were the following: 120g weight, released from a height of 40cm leading to a maximum skull displacement of 1,5mm (Chandrasekar et al., 2019). Immediately after the impact, mice received 100% O_2_ and the temporary apnea time measured. The mice were kept under anesthesia (4% Servofluran, 96% O_2_) while the skin was sutured using Prolene 6.0 surgical thread. Mice received additional opiate (0.05mg/kg buprenorphine) treatment for the next 24h, access to soft food pellets and were checked for their well being using a score sheet. The Neurological Severity Score (NSS) was assessed after 3h, 1 dpi and at 7 dpi and never exceeded the score 1 for any mouse; as such, no animal met the criteria for early sacrifice. Sham operated mice underwent the same treatment except for the impact injury.

### Immunohistochemistry

Brain samples were processed as previously described (olde Heuvel et al., 2019). Briefly, mice were sacrificed by trans-cardial perfusion with 4% PFA in PBS, and brains were dissected and postfixed in 4% PFA overnight. Brains were then washed and transferred to 30% Sucrose for 2 days, after which the samples were embedded in OCT (Tissue Tek, Sakura, Germany. 40 micron sections were cut with a cryostat (Leica CM 1950 AG Protect cryostat). Sections spanning the injury site were selected and blocked (3% BSA, 0.3% Triton in 1x PBS) for 2h at room temperature, followed by incubation for 48h at 4°C with primary antibodies diluted in blocking buffer. Sections were washed 3x 30 min with PBS, and incubated for 2h at RT with secondary antibodies diluted in blocking buffer. The sections were washed 3x 30 min with PBS and mounted using Prolong Gold Antifade Mounting Medium (Invitrogen, Germany). The full list of antibodies used can be found in Supp. Table 2.

### Single molecule in situ hybridization

Single mRNA fluorescent in situ hybridisation (Wang et al., 2012) was performed according to manufacturer instructions (ACDBio, RNAscope, fluorescent In Situ Hybridisation) for fixed frozen tissue sections, all reagents and probes were provided by ACDBio; Supp. Table 3) with small modifications (olde Heuvel et al., 2019). Briefly, cryosections were mounted in superfrost plus glass slides and frozen at −80°C for at least 24h before the actual procedure. Sections were retrieved from the freezer and kept at RT before being washed in PBS for 5 min at RT. An antigen retrieval step was performed at 100°C for 5 min, followed by washing twice in dH_2_0 and once in ethanol. The slides were then pretreated with reagent III for 30 min at 40°C, and then washed twice in dH_2_0. Probe hybridisation (OPG and VGLUT2) was performed for 4.5h at 40°C, followed by washing 2x 2 min in washing buffer. First amplification was performed by incubating the slides in amplification-1 reagent for 30 min at 40°C, followed by washing 2x 2 min in washing buffer. Second amplification was performed by incubating the slides in amplification-2 reagent for 15 min at 40°C, followed by washing 2x 2 min in washing buffer. The last amplification was performed by incubating amplification-3 reagent for 30 min at 40°C, followed by washing 2x 2 min in washing buffer. The detection step was performed by incubating amplification-4 reagent for 45 min at 40°C, followed by washing 2x 10 min in washing buffer. Co-immunostaining was performed directly after the final detection, whereby the slides were blocked for 1 h in blocking buffer (% BSA, 0.3% triton in 1x PBS). Primary antibodies for GFP (Chicken; 1:500; Abcam Ab13970), NeuN (Guinea Pig; 1:250; SySy 266004) and RFP (Camel; 1:250; Nanotech N0404-AT565-L) were diluted in blocking buffer and incubated overnight at 4°C, followed by 3x 30 min washing in PBS. Secondary antibodies for NeuN (Goat Anti-guinea pig IgG H&L Alexa Fluor 405; 1:250; abcam ab175678) and GFP (Alexa Fluor 488 AffiniPure Donkey Anti-Chicken IgY (IgG) (H+L); 1:250; Jackson 703-545-155) were diluted in blocking buffer and incubated for 2 h at RT, followed by 3x 30 min washing in PBS. The sections were mounted using fluorogold ProLong antifade mounting medium (Invitrogen).

### Image Acquisition and analysis

Confocal images were acquired in 1024 x 1024 pixel and 12-bit format, with a Leica DMi8 inverted microscope, equipped with an ACS APO 40x oil objective. Parameters were set to obtain the signals from the stained antibody or mRNA and at the same time avoiding saturation. All fluorescent channels were acquired independently, to avoid cross-bleed. 3-4 sections spanning the core and perilesional area of the impact site were imaged of each mouse. Optical stacks of 30 micron were obtained and imported to ImageJ. For image analysis, stacks were collapsed in maximum intensity projection pictures and mean gray value or cell density per fixed region of interest (ROI) was measured.

CREB phosphorylation was visualised by acquiring a mosaic image of 2×1 optical stacks covering the center of the injection / injury site in layer II/III of the cerebral cortex. After importing the images to Image J a 350μm x 200μm area centered on the injury site was considered as ROI and mean gray value of pCREB was measured in every NeuN+ or IBA1+ nucleus. To analyze the cell density mosaic images corresponding to 3×1 optical stacks spanning the core and perilesional area in the II/III cortical layer were acquired. After importing the images to ImageJ 250μm x 200μm ROIs were selected for analysis as indicated in Supp. Fig. 2A and the cells within the ROIs were quantified. For OPG mRNA quantification a 2×2 mosaic image located in the core area of the injury in layer II/III was acquired. The images were imported to Image J and after maximum intensity projection, OPG mRNA density of each VGlut2+ Neuron was quantified.

Synaptic density was detected after producing a mosaic image corresponding to 6 x 6 single optical sections (acquired with a 63x oil objective) with 1 μm thickness. Each cortical section was imaged at a fixed depth of 10 μm inside the section, the composite image was positioned so that an uninterrupted coverage of the impact site with perilesional area was acquired. Quantification of density of synapses was done with the IMARIS software (Bitplane AG, Zurich, CH), ROI were positioned at fixed distance into the cerebral cortex (layer II/III) at the core of the injury and at the fixed distance from the core (150 μm) at the perilesional area. The number of SHANK+ synapses per unit area was counted using IMARIS as parameters; estimated XY diameter of 0.7 μm, Quality of 28% and a region threshold of 0. The parameters were kept constant for each ROI.

### Nanostring targeted transcriptome and bioinformatic analysis

At 3h post TBI mice were euthanized, the brain quickly extracted. The cortex was dissected in ice cold PBS and snap frozen on dry ice. Cortical tissue was stored at −80°C until further use. RNA was extracted using the ISOLATE II RNA/DNA/Protein kit (Bioline, Germany) following the manufacturer’s instructions. Reagents and buffers were all provided by the company. Briefly, cortical samples were homogenized in 300 μl lysis buffer, loaded on a DNA column and centrifuged at 14,000g for 1 min. The flow-through was collected and adjusted with ethanol before loading it on the RNA column and centrifuged at 3,500g for 1 min. The column was washed 3x by adding 400 μl of washing buffer and centrifuging at 14,000g for 1 min, followed by a dry spin at 14,000g for 2 min. The RNA was eluted by adding 50 μl of elution buffer and centrifuging at 200g for 2 min followed by a final dry spin at 14,000g for 1 min. The isolated RNA was collected and stored at −80°C until further use. The RNA concentration and quality were determined with the Nanodrop 2000 and samples were sent to Proteros (Germany) for gene expression profiling using NanoString technology.

The raw data files were loaded in R software and the dataset for each sample was preliminary subjected to quality control assessment (QCA). Normalized data were then subjected to principal component analysis (PCA) to display group-based clustering. Confidence ellipses (assuming multivariate normal distribution) with the first two principal components were plotted to validate further analysis. Modified linear modeling-based analysis was then applied to the data to identify RTK showing a significant increase or decrease in expression at different conditions.

### Neurological Severity score and Whisking analysis

Neurological severity score (NSS) and whisking analysis were performed at 1d, 3d and 7d post TBI. NSS wwas performed as previously reported (Flierl et al., 2009). Briefly, mice were subjected to ten tasks to investigate neurological damage after TBI. The tasks included; exit circle test, seeking behaviour, showing mono-/hemiparesis, straight walk test, startle reflex test, beam balance test, round stick balance test and beam walk test (1, 2 and 3 cm). If the mouse failed the test, a point was awarded.

Whisking analyses were performed as previously reported (Wanner et al., 2020). Briefly, mice were held and fixed in one place, in such a way that the head and whiskers could move freely. Whiskers movement was videotaped for 50 sec by a highspeed camera (Basler acA1300-60gc) at 100 Hz. General whisking activity was analyzed by measuring the total amount of whisking events per video and 1 second fragments were processed and analyzed with the Templo and Vicon Motus 2D software (CONTEMPLAS, Germany). Amplitude and velocity of the whiskers were assessed and calculated.

### Clinical cohort and ELISA assay for OPG measurements

The collection of CSF samples from human TBI patients was authorized by The Alfred Hospital Ethical Committee (Melbourne) no. 194-05 to Cristina Morganti-Kossmann and by the Ulm Universtiy Ethical Committee no. 12/19-22050. Clinical and demographic characteristics of the patients are reported in Table 1. Patients for this cohort were recruited at the Alfred Hospital, Melbourne; informed consent was obtained from the next of the kin. Inclusion criteria were: severe TBI with a post-resuscitation GCS ≤8 (unless initial GCS>9 was followed by deterioration requiring intubation) and, upon CT imaging, the need for an extraventricular drain (EVD) for ICP monitoring and therapeutic drainage of CSF. CSF was collected over 24⍰h and kept at 4°C; samples were obtained on admission (day 0) and daily up to day 7 after injury. Within an hour from collection, samples were centrifuged at 2000g for 15 min at 4°C and stored at −80°C until analysis. Exclusion criteria comprised pregnancy, neurodegenerative diseases, HIV and other chronic infection/inflammatory diseases, or history of TBI. Out of the 42 TBI patients constituting the original cohort (Yan et al., 2014), we selected a total of 67 samples corresponding to D0, D1, D3 and D7 after injury.

OPG in the CSF was measured using the Human TNFRSF11B(OPG) Elisa kit (Thermo Fisher Scientific, Germany). The assay was performed according to the manufacturer’s instructions. 100 μL of diluted sample (1:1 in the sample diluent) was used. Absorbance at 450 nm was measured in ELx800 Microplate Reader (BioTek) and concentrations were calculated according to the standard curve.

### Statistics

Statistical analysis was performed using GraphPad Prism version 8 software (GraphPad software, USA). The Shapiro-Wilk test was performed to test groups for normality. Grouped analysis was performed with the mean of each animal and using two-way ANOVA (analysis of variance) with Tukey’s multiple correction. Depending on normality, grouped analysis for the ELISA data was performed using ANOVA with Tukey’s multiple correction or Kruskal-Wallis test with Dunn’s multiple correction. Data are depicted as histograms with mean ± SD or scatterplot with mean ± SD. Statistical significance was set at p < 0.05.

## Acknowledgement

This work has been supported by the Deutsche Forschungsgemeinschaft as part of the Collaborative Research Center 1149 “Danger Response, Disturbance Factors and Regenerative Potential after Acute Trauma” (DFG No. 251293561). FR is also supported by DFG through the individual grants no. 443642953, no. 431995586 and no. 446067541, and by the ERANET-NEURON initiative “External Insults to the Nervous System” as part of the MICRONET consortium (funded by BMBF: FKZ 01EW1705A). SL is supported by the China Scholarship Council (No. 201908320357). We are grateful to Prof. Stefan Just and to Prof. Anita Ignatius for the use of the confocal microscope and the histology laboratory. We would like to thank the colleagues of the German Center for Neurodegenerative Disease (DZNE)-Ulm for advice and discussion. Technical support by Thomas Lenk and Gizem Yartas was highly appreciated.

## Authors contribution

FR designed and conceived the study. AF, FoH, RR, ZL performed the in vivo experiments. RR performed the bioinformatic analysis of the nanostring data. SSK performed the ELISA and the SIMOA assays on CSF samples. AC and CMK contributed the human CSF samples from the Melbourne cohort. DB prepared the AAV vectors used in the study. AHH and HB cloned the nuclear calcium buffers. MHL contributed to the establishment of the TBI model. BK contributed to the analysis of the whisking behaviour. FR, AF, FoH SL, AL, TB, MHL and BK cooperated in the critical analysis of the data and in the preparation of the initial draft. FR, CMK and HB contributed to the final version of the manuscript.

---

***Supplementary Figure 1 - Glial CREB phosphorylation post injury is limited to IBA1+ cells.***

A) Confocal images of somatosensory cortex layer 2 and 3 showing NeuN (magenta) and RFP (red). Scale bar: 5Oμm.

B) AAV9 viruses used in these experiments show an infection rate of 89.98 ± 14.47%. Data is shown as mean ± SD. N=8.

C) Confocal images of somatosensory cortex layer 2-3 showing pCREB (cyan), IBA1 (magenta) and GFAP (magenta). Scale Bar: 5Oμm.

D) 97.76 ± 0.83% of pCREB+ glial cells are IBA1+ while 1.24 ± 0.83% are GFAP+. Data is shown as mean ± SD. N=4.

***Supplementary Figure 2 - Blunting neuronal nuclear calcium signaling did not alter IBA1+/TMEM119+ cell density and percentage in the perilesional area post TBI.***

A) Confocal image of layer 2-3 showcasing the positioning of ROIs for the core region (centered on the Injury axis) and the perilesional area (200μm away from the axis).

B) No significant difference in IBA1+ and IBA1+/TMEM119+ cell density in the perilesional area 24h post injury compared to sham or nuclear calcium buffering (CS= 4.10 ± 0.47, CT = 6.73 ± 1.62 and PT = 8.92 ± 1.84 for IBA1+; CS = 4 ± 0.48, CT = 6.44 ± 1.49 and PT = 8.71 ± 1.79 for IBA1+/TMEM119+). Significant increase in IBA1+/CD11c+ cell density in the perilesional area compared to sham but not after nuclear calcium buffering (CS vs CT 0.5 ± 0.39 vs 5.5 ± 1.77; CT vs PT 5.5 ± 1.77 vs 5.5 ± 0.97). Data is shown as mean ± SD. N=4. *p<0.05, **p<0.01.

C) No significant differences in the percentage of TMEM119+ cells can be observed 24h post injury between all groups (CS = 97.46 ± 2.34%; CT = 95.91 ± 2.5%; PS = 96.39 ± 1.78%; PT = 97.69 ± 0.87%). Significant increase in CD11+ expression 24h after injury compared to sham but not to PV.NLS TBI (CS vs CT 12.17 ± 10.2% vs 80.3 ± 8.32%; CT vs PT 80.3 ± 8.32% vs 64.44 ± 20.57%). Data shown as mean ± SD. N=4. **p<0.01, ***p<0.001.

***Supplementary Figure 3 - Blunting neuronal nuclear calcium signaling did not alter amplitude and velocity of whisker movement post TBI.***

A) Amplitudes of the affected whisker are not significantly changed 24h after injury in relation to D-1 after Control sham (CS), Control TBI (CT), PV.NLS sham (PS) and PV.NLS TBI (PT) treatment (CS = 0.88 ± 0.22; CT = 1.1 ± 0.19; PS = 0.93 ± 0.42; PT = 1.269 ± 0.56). Data shown as mean ± SD. N=7.

B) Amplitudes of the unaffected whisker are not significantly changed 24h after injury in relation to D-1 after CS, CT, PS and PT treatment (CS = 1.07 ± 0.39; CT = 1.05 ± 0.35; PS = 1.14 ± 0.45; PT = 1.22 ± 0.45). Data shown as mean ± SD. N=7.

C) The ratios between the amplitudes of the affected and unaffected whiskers are not significantly changed 24h after injury in relation to D-1 after CS, CT, PS and PT treatment (CS = 0.77 ± 0.27; CT = 1.24 ± 0.42; PS = 0.91 ± 0.45; PT = 1.10 ± 0.55). Data shown as mean ± SD. N=7.

D) Velocities of the affected whisker are not significantly changed 24h after injury in relation to D-1 after CS, CT, PS and PT treatment (CS = 1.1 ± 0.38; CT = 1.02 ± 0.29; PS = 0.95 ± 0.43; PT = 1.43 ± 0.78). Data shown as mean ± SD. N=7.

E) Velocities of the unaffected whisker are not significantly changed 24h after injury in relation to D-1 after CS, CT, PS and PT treatment (CS = 0.95 ± 0.66; CT = 0.82 ± 0.26; PS = 1.33 ± 0.75; PT = 1.2 ± 0.62). Data shown as mean ± SD. N=7.

F) The ratio between the velocities of the affected and unaffected whiskers are not significantly changed 24h after injury in relation to D-1 after CS, CT, PS and PT treatment (CS = 1.74 ± 1.26; CT = 1.27 ± 0.72; PS = 0.89 ± 0.54; PT = 0.75 ± 0.33). Data shown as mean ± SD. N=7.

***Supplementary Figure 4 - Re-expression of osteoprotegerin together with nuclear calcium buffering massively induces neuronal OPG mRNA intensity.***

A) Confocal images of somatosensory cortex layer 2-3 obtained from Control sham (CS), Control TBI (CT), PV.NLS TBI (PT) and PV.NLS OPG TBI (POT) mice. Shown are DAPI (blue), RFP (red) and OPG mRNA (white). Scale bar overview: 2Oμm.

B) Significant increase in neuronal OPG mRNA intensity (CS vs CT; 181.8 ± 24.34 vs 224.3 ± 35.87). Buffering of nuclear calcium signaling decreased neuronal OPG mRNA intensity (CT vs PT; 224.3 ± 35.87 vs 184.1 ± 24.11). Re-expression of OPG together with nuclear calcium buffering increased the neuronal OPG mRNA intensity massively (PT vs POT; 184.1 ± 24.11 vs 1386 ± 435). Data is shown as mean ± SD. N=4.

***Supplementary Table 1 - Targeted transcriptome analysis reveals new neuronal nuclear calcium-regulated mediators of neuro-glia crosstalk upon TBI.***

All significantly differentially regulated genes between the PV.NLS TBI (PT) and Control TBI (CT) /PV.NLS Sham (PS) are listed with name, log FC and their adjusted p-value.

***Supplementary Table 2 - Antibodies and Constructs.***

Full information on antibodies and constructs has been listed.

## References

Ahlgren H, Bas-Orth C, Freitag HE, Hellwig A, Ottersen OP, Bading H. The nuclear calcium signaling target, activating transcription factor 3 (ATF3), protects against dendrotoxicity and facilitates the recovery of synaptic transmission after an excitotoxic insult. J Biol Chem. 2014 Apr 4;289(14):9970–82. doi: 10.1074/jbc.M113.502914.

Allitt BJ, Iva P, Yan EB, Rajan R. Hypo-excitation across all cortical laminae defines intermediate stages of cortical neuronal dysfunction in diffuse traumatic brain injury. Neuroscience. 2016 Oct 15;334:290–308. doi: 10.1016/j.neuroscience.2016.08.018.

Ashcroft AJ, Cruickshank SM, Croucher PI, Perry MJ, Rollinson S, Lippitt JM, Child JA, Dunstan C, Felsburg PJ, Morgan GJ, Carding SR. Colonic dendritic cells, intestinal inflammation, and T cell-mediated bone destruction are modulated by recombinant osteoprotegerin. Immunity. 2003 Dec;19(6):849–61. doi: 10.1016/s1074-7613(03)00326-1.

Bading H. Nuclear calcium signalling in the regulation of brain function. Nat Rev Neurosci. 2013 Sep;14(9):593–608. doi: 10.1038/nrn3531.

Bading H. Therapeutic targeting of the pathological triad of extrasynaptic NMDA receptor signaling in neurodegenerations. J Exp Med. 2017 Mar 6;214(3):569–578. doi: 10.1084/jem.20161673.

Bas-Orth C, Tan YW, Lau D, Bading H. Synaptic Activity Drives a Genomic Program That Promotes a Neuronal Warburg Effect. J Biol Chem. 2017 Mar 31;292(13):5183–5194. doi: 10.1074/jbc.M116.761106.

Bell KF, Bent RJ, Meese-Tamuri S, Ali A, Forder JP, Aarts MM. Calmodulin kinase IV-dependent CREB activation is required for neuroprotection via NMDA receptor-PSD95 disruption. J Neurochem. 2013 Jul;126(2):274–87. doi: 10.1111/jnc.12176.

Biber K, Neumann H, Inoue K, Boddeke HW. Neuronal ‘On’ and ‘Off signals control microglia. Trends Neurosci. 2007 Nov;30(11):596–602. doi: 10.1016/j.tins.2007.08.007.

Bogie JF, Boelen E, Louagie E, Delputte P, Elewaut D, van Horssen J, Hendriks JJ, Hellings N. CD169 is a marker for highly pathogenic phagocytes in multiple sclerosis. Mult Scler. 2018 Mar;24(3):290–300. doi: 10.1177/1352458517698759.

Buchthal B, Lau D, Weiss U, Weislogel JM, Bading H. Nuclear calcium signaling controls methyl-CpG-binding protein 2 (MeCP2) phosphorylation on serine 421 following synaptic activity. J Biol Chem. 2012 Sep 7;287(37):30967–74. doi: 10.1074/jbc.M112.382507.

Carron SF, Alwis DS, Rajan R. Traumatic Brain Injury and Neuronal Functionality Changes in Sensory Cortex. Front Syst Neurosci. 2016 Jun 2;10:47. doi: 10.3389/fnsys.2016.00047.

Chagraoui H, Tulliez M, Smayra T, Komura E, Giraudier S, Yun T, Lassau N, Vainchenker W, Wendling F. Stimulation of osteoprotegerin production is responsible for osteosclerosis in mice overexpressing TPO. Blood. 2003 Apr 15;101(8):2983–9. doi: 10.1182/blood-2002-09-2839.

Chandrasekar A, Heuvel FO, Tar L, Hagenston AM, Palmer A, Linkus B, Ludolph AC, Huber-Lang M, Boeckers T, Bading H, Roselli F. Parvalbumin Interneurons Shape Neuronal Vulnerability in Blunt TBI. Cereb Cortex. 2019 Jun 1;29(6):2701–2715. doi: 10.1093/cercor/bhy139.

Chandrasekar A, Olde Heuvel F, Wepler M, Rehman R, Palmer A, Catanese A, Linkus B, Ludolph A, Boeckers T, Huber-Lang M, Radermacher P, Roselli F. The Neuroprotective Effect of Ethanol Intoxication in Traumatic Brain Injury Is Associated with the Suppression of ErbB Signaling in Parvalbumin-Positive Interneurons. J Neurotrauma. 2018 Nov 15;35(22):2718–2735. doi: 10.1089/neu.2017.5270.

Commisso B, Ding L, Varadi K, Gorges M, Bayer D, Boeckers TM, Ludolph AC, Kassubek J, Müller OJ, Roselli F. Stage-dependent remodeling of projections to motor cortex in ALS mouse model revealed by a new variant retrograde-AAV9. Elife. 2018 Aug 23;7:e36892. doi: 10.7554/eLife.36892.

Chino T, Draves KE, Clark EA. Regulation of dendritic cell survival and cytokine production by osteoprotegerin. J Leukoc Biol. 2009 Oct;86(4):933–40. doi: 10.1189/jlb.0708419.

Depp C, Bas-Orth C, Schroeder L, Hellwig A, Bading H. Synaptic Activity Protects Neurons Against Calcium-Mediated Oxidation and Contraction of Mitochondria During Excitotoxicity. Antioxid Redox Signal. 2018 Oct 20;29(12):1109–1124. doi: 10.1089/ars.2017.7092.

Di Liberto G, Pantelyushin S, Kreutzfeldt M, Page N, Musardo S, Coras R, Steinbach K, Vincenti I, Klimek B, Lingner T, Salinas G, Lin-Marq N, Staszewski O, Costa Jordão MJ, Wagner I, Egervari K, Mack M, Bellone C, Blümcke I, Prinz M, Pinschewer DD, Merkler D. Neurons under T Cell Attack Coordinate Phagocyte-Mediated Synaptic Stripping. Cell. 2018 Oct 4;175(2):458–471.e19. doi: 10.1016/j.cell.2018.07.049.

Flierl MA, Stahel PF, Beauchamp KM, Morgan SJ, Smith WR, Shohami E. Mouse closed head injury model induced by a weight-drop device. Nat Protoc. 2009;4(9):1328–37. doi: 10.1038/nprot.2009.148.

Förstner P, Knöll B. Interference of neuronal activity-mediated gene expression through serum response factor deletion enhances mortality and hyperactivity after traumatic brain injury. FASEB J. 2020 Mar;34(3):3855–3873. doi: 10.1096/fj.201902257RR.

Förstner P, Rehman R, Anastasiadou S, Haffner-Luntzer M, Sinske D, Ignatius A, Roselli F, Knöll B. Neuroinflammation after Traumatic Brain Injury Is Enhanced in Activating Transcription Factor 3 Mutant Mice. J Neurotrauma. 2018 Oct 1;35(19):2317–2329. doi: 10.1089/neu.2017.5593.

Glasnović A, O’Mara N, Kovačić N, Grčević D, Gajović S. RANK/RANKL/OPG Signaling in the Brain: A Systematic Review of the Literature. Front Neurol. 2020 Nov 19;11:590480. doi: 10.3389/fneur.2020.590480.

Glasnović A, Stojić M, Dežmalj L, Tudorić-Đeno I, Romić D, Jeleč V, Vrca A, Vuletić V, Grčević D. RANKL/RANK/OPG Axis Is Deregulated in the Cerebrospinal Fluid of Multiple Sclerosis Patients at Clinical Onset. Neuroimmunomodulation. 2018;25(1):23–33. doi: 10.1159/000488988.

Guerrini MM, Okamoto K, Komatsu N, Sawa S, Danks L, Penninger JM, Nakashima T, Takayanagi H. Inhibition of the TNF Family Cytokine RANKL Prevents Autoimmune Inflammation in the Central Nervous System. Immunity. 2015 Dec 15;43(6):1174–85. doi: 10.1016/j.immuni.2015.10.017.

Hammond JW, Bellizzi MJ, Ware C, Qiu WQ, Saminathan P, Li H, Luo S, Ma SA, Li Y, Gelbard HA. Complement-dependent synapse loss and microgliosis in a mouse model of multiple sclerosis. Brain Behav Immun. 2020 Jul;87:739–750. doi: 10.1016/j.bbi.2020.03.004.

Hanada R. The role of the RANKL/RANK/OPG system in the central nervous systems (CNS). J Bone Miner Metab. 2021 Jan;39(1):64–70. doi: 10.1007/s00774-020-01143-9.

Hardingham GE, Chawla S, Johnson CM, Bading H. Distinct functions of nuclear and cytoplasmic calcium in the control of gene expression. Nature. 1997 Jan 16;385(6613):260–5. doi: 10.1038/385260a0.

Hoek RM, Ruuls SR, Murphy CA, Wright GJ, Goddard R, Zurawski SM, Blom B, Homola ME, Streit WJ, Brown MH, Barclay AN, Sedgwick JD. Down-regulation of the macrophage lineage through interaction with OX2 (CD200). Science. 2000 Dec 1;290(5497):1768–71. doi: 10.1126/science.290.5497.1768.

Hofbauer LC, Cepok S, Hemmer B. Osteoprotegerin is highly expressed in the spinal cord and cerebrospinal fluid. Acta Neuropathol. 2004 Jun;107(6):575–7, author reply 578. doi: 10.1007/s00401-004-0854-y.

Jassam YN, Izzy S, Whalen M, McGavern DB, El Khoury J. Neuroimmunology of Traumatic Brain Injury: Time for a Paradigm Shift. Neuron. 2017 Sep 13;95(6):1246–1265. doi: 10.1016/j.neuron.2017.07.010.

Jensen JK, Ueland T, Atar D, Gullestad L, Mickley H, Aukrust P, Januzzi JL. Osteoprotegerin concentrations and prognosis in acute ischaemic stroke. J Intern Med. 2010 Apr;267(4):410–7. doi: 10.1111/j.1365-2796.2009.02163.x.

Johnstone VP, Shultz SR, Yan EB, O’Brien TJ, Rajan R. The acute phase of mild traumatic brain injury is characterized by a distance-dependent neuronal hypoactivity. J Neurotrauma. 2014 Nov 15;31(22):1881–95. doi: 10.1089/neu.2014.3343.

Keren-Shaul H, Spinrad A, Weiner A, Matcovitch-Natan O, Dvir-Szternfeld R, Ulland TK, David E, Baruch K, Lara-Astaiso D, Toth B, Itzkovitz S, Colonna M, Schwartz M, Amit I. A Unique Microglia Type Associated with Restricting Development of Alzheimer’s Disease. Cell. 2017 Jun 15;169(7):1276–1290.e17. doi: 10.1016/j.cell.2017.05.018.

Kichev A, Eede P, Gressens P, Thornton C, Hagberg H. Implicating Receptor Activator of NF-κB (RANK)/RANK Ligand Signalling in Microglial Responses to Toll-Like Receptor Stimuli. Dev Neurosci. 2017;39(1-4):192–206. doi: 10.1159/000464244.

Kichev A, Rousset CI, Baburamani AA, Levison SW, Wood TL, Gressens P, Thornton C, Hagberg H. Tumor necrosis factor-related apoptosis-inducing ligand (TRAIL) signaling and cell death in the immature central nervous system after hypoxia-ischemia and inflammation. J Biol Chem. 2014 Mar 28;289(13):9430–9. doi: 10.1074/jbc.M113.512350.

Krasemann S, Madore C, Cialic R, Baufeld C, Calcagno N, El Fatimy R, Beckers L, O’Loughlin E, Xu Y, Fanek Z, Greco DJ, Smith ST, Tweet G, Humulock Z, Zrzavy T, Conde-Sanroman P, Gacias M, Weng Z, Chen H, Tjon E, Mazaheri F, Hartmann K, Madi A, Ulrich JD, Glatzel M, Worthmann A, Heeren J, Budnik B, Lemere C, Ikezu T, Heppner FL, Litvak V, Holtzman DM, Lassmann H, Weiner HL, Ochando J, Haass C, Butovsky O. The TREM2-APOE Pathway Drives the Transcriptional Phenotype of Dysfunctional Microglia in Neurodegenerative Diseases. Immunity. 2017 Sep 19;47(3):566–581.e9. doi: 10.1016/j.immuni.2017.08.008.

Lago N, Pannunzio B, Amo-Aparicio J, López-Vales R, Peluffo H. CD200 modulates spinal cord injury neuroinflammation and outcome through CD200R1. Brain Behav Immun. 2018 Oct;73:416–426. doi: 10.1016/j.bbi.2018.06.002.

Li B, Jie W, Huang L, Wei P, Li S, Luo Z, Friedman AK, Meredith AL, Han MH, Zhu XH, Gao TM. Nuclear BK channels regulate gene expression via the control of nuclear calcium signaling. Nat Neurosci. 2014 Aug;17(8):1055–63. doi: 10.1038/nn.3744.

Lim TK, Ruthazer ES. Microglial trogocytosis and the complement system regulate axonal pruning in vivo. Elife. 2021 Mar 16;10:e62167. doi: 10.7554/eLife.62167.

Matyas F, Sreenivasan V, Marbach F, Wacongne C, Barsy B, Mateo C, Aronoff R, Petersen CC. Motor control by sensory cortex. Science. 2010 Nov 26;330(6008):1240–3. doi: 10.1126/science.1195797.

Mauceri D, Hagenston AM, Schramm K, Weiss U, Bading H. Nuclear Calcium Buffering Capacity Shapes Neuronal Architecture. J Biol Chem. 2015 Sep 18;290(38):23039–49. doi: 10.1074/jbc.M115.654962.

McGinn MJ, Kelley BJ, Akinyi L, Oli MW, Liu MC, Hayes RL, Wang KK, Povlishock JT. Biochemical, structural, and biomarker evidence for calpain-mediated cytoskeletal change after diffuse brain injury uncomplicated by contusion. J Neuropathol Exp Neurol. 2009 Mar;68(3):241–9. doi: 10.1097/NEN.0b013e3181996bfe.

Morganti-Kossmann MC, Semple BD, Hellewell SC, Bye N, Ziebell JM. The complexity of neuroinflammation consequent to traumatic brain injury: from research evidence to potential treatments. Acta Neuropathol. 2019 May;137(5):731–755. doi: 10.1007/s00401-018-1944-6. Epub 2018 Dec 7.

Mozolewski P, Jeziorek M, Schuster CM, Bading H, Frost B, Dobrowolski R. The role of nuclear Ca2+ in maintaining neuronal homeostasis and brain health. J Cell Sci. 2021 Apr 15;134(8):jcs254904. doi: 10.1242/jcs.254904.

Onodera J, Nagata H, Nakashima A, Ikegaya Y, Koyama R. Neuronal brain-derived neurotrophic factor manipulates microglial dynamics. Glia. 2021 Apr;69(4):890–904. doi: 10.1002/glia.23934.

Olde Heuvel F, Holl S, Chandrasekar A, Li Z, Wang Y, Rehman R, Förstner P, Sinske D, Palmer A, Wiesner D, Ludolph A, Huber-Lang M, Relja B, Wirth T, Röszer T, Baumann B, Boeckers T, Knöll B, Roselli F. STAT6 mediates the effect of ethanol on neuroinflammatory response in TBI. Brain Behav Immun. 2019 Oct;81:228–246. doi: 10.1016/j.bbi.2019.06.019.

Ouali Alami N, Schurr C, Olde Heuvel F, Tang L, Li Q, Tasdogan A, Kimbara A, Nettekoven M, Ottaviani G, Raposo C, Röver S, Rogers-Evans M, Rothenhäusler B, Ullmer C, Fingerle J, Grether U, Knuesel I, Boeckers TM, Ludolph A, Wirth T, Roselli F, Baumann B. NF-κB activation in astrocytes drives a stagespecific beneficial neuroimmunological response in ALS. EMBO J. 2018 Aug 15;37(16):e98697. doi: 10.15252/embj.201798697.

Piao CS, Stoica BA, Wu J, Sabirzhanov B, Zhao Z, Cabatbat R, Loane DJ, Faden AI. Late exercise reduces neuroinflammation and cognitive dysfunction after traumatic brain injury. Neurobiol Dis. 2013 Jun;54:252–63. doi: 10.1016/j.nbd.2012.12.017.

Pohl D, Bittigau P, Ishimaru MJ, Stadthaus D, Hübner C, Olney JW, Turski L, Ikonomidou C. N-Methyl-D-aspartate antagonists and apoptotic cell death triggered by head trauma in developing rat brain. Proc Natl Acad Sci U S A. 1999 Mar 2;96(5):2508–13. doi: 10.1073/pnas.96.5.2508.

Pruunsild P, Bengtson CP, Bading H. Networks of Cultured iPSC-Derived Neurons Reveal the Human Synaptic Activity-Regulated Adaptive Gene Program. Cell Rep. 2017 Jan 3;18(1):122–135. doi: 10.1016/j.celrep.2016.12.018.

Pusl T, Wu JJ, Zimmerman TL, Zhang L, Ehrlich BE, Berchtold MW, Hoek JB, Karpen SJ, Nathanson MH, Bennett AM. Epidermal growth factor-mediated activation of the ETS domain transcription factor Elk-1 requires nuclear calcium. J Biol Chem. 2002 Jul 26;277(30):27517–27. doi: 10.1074/jbc.M203002200.

Rajan WD, Wojtas B, Gielniewski B, Miró-Mur F, Pedragosa J, Zawadzka M, Pilanc P, Planas AM, Kaminska B. Defining molecular identity and fates of CNS-border associated macrophages after ischemic stroke in rodents and humans. Neurobiol Dis. 2020 Apr;137:104722. doi: 10.1016/j.nbd.2019.104722.

Rehman R, Tar L, Olamide AJ, Li Z, Kassubek J, Böckers T, Weishaupt J, Ludolph A, Wiesner D, Roselli F. Acute TBK1/IKK-ε Inhibition Enhances the Generation of Disease-Associated Microglia-Like Phenotype Upon Cortical Stab-Wound Injury. Front Aging Neurosci. 2021 Jul 13;13:684171. doi: 10.3389/fnagi.2021.684171.

Robertson CL. Mitochondrial dysfunction contributes to cell death following traumatic brain injury in adult and immature animals. J Bioenerg Biomembr. 2004 Aug;36(4):363–8. doi: 10.1023/B:JOBB.0000041769.06954.e4.

Roselli F, Caroni P. From intrinsic firing properties to selective neuronal vulnerability in neurodegenerative diseases. Neuron. 2015 Mar 4;85(5):901–10. doi: 10.1016/j.neuron.2014.12.063.

Russo MV, McGavern DB. Inflammatory neuroprotection following traumatic brain injury. Science. 2016 Aug 19;353(6301):783–5. doi: 10.1126/science.aaf6260.

Shimamura M, Nakagami H, Osako MK, Kurinami H, Koriyama H, Zhengda P, Tomioka H, Tenma A, Wakayama K, Morishita R. OPG/RANKL/RANK axis is a critical inflammatory signaling system in ischemic brain in mice. Proc Natl Acad Sci U S A. 2014 Jun 3;111(22):8191–6. doi: 10.1073/pnas.1400544111.

Schlumm F, Mauceri D, Freitag HE, Bading H. Nuclear calcium signaling regulates nuclear export of a subset of class IIa histone deacetylases following synaptic activity. J Biol Chem. 2013 Mar 22;288(12):8074–8084. doi: 10.1074/jbc.M112.432773.

Shaw JA, Perry VH, Mellanby J. MHC class II expression by microglia in tetanus toxin-induced experimental epilepsy in the rat. Neuropathol Appl Neurobiol. 1994 Aug;20(4):392–8. doi: 10.1111/j.1365-2990.1994.tb00985.x.

Simonetti M, Hagenston AM, Vardeh D, Freitag HE, Mauceri D, Lu J, Satagopam VP, Schneider R, Costigan M, Bading H, Kuner R. Nuclear calcium signaling in spinal neurons drives a genomic program required for persistent inflammatory pain. Neuron. 2013 Jan 9;77(1):43–57. doi: 10.1016/j.neuron.2012.10.037.

Sreenivasan V, Esmaeili V, Kiritani T, Galan K, Crochet S, Petersen CCH. Movement Initiation Signals in Mouse Whisker Motor Cortex. Neuron. 2016 Dec 21;92(6):1368–1382. doi: 10.1016/j.neuron.2016.12.001.

Stover JF, Morganti-Kosmann MC, Lenzlinger PM, Stocker R, Kempski OS, Kossmann T. Glutamate and taurine are increased in ventricular cerebrospinal fluid of severely brain-injured patients. J Neurotrauma. 1999 Feb;16(2):135–42. doi: 10.1089/neu.1999.16.135.

Szepesi Z, Manouchehrian O, Bachiller S, Deierborg T. Bidirectional Microglia-Neuron Communication in Health and Disease. Front Cell Neurosci. 2018 Sep 27;12:323. doi: 10.3389/fncel.2018.00323.

Tamakoshi K, Hayao K, Takahashi H. Early Exercise after Intracerebral Hemorrhage Inhibits Inflammation and Promotes Neuroprotection in the Sensorimotor Cortex in Rats. Neuroscience. 2020 Jul 1;438:86–99. doi: 10.1016/j.neuroscience.2020.05.003.

Umpierre AD, Bystrom LL, Ying Y, Liu YU, Worrell G, Wu LJ. Microglial calcium signaling is attuned to neuronal activity in awake mice. Elife. 2020 Jul 27;9:e56502. doi: 10.7554/eLife.56502.

Vasek MJ, Garber C, Dorsey D, Durrant DM, Bollman B, Soung A, Yu J, Perez-Torres C, Frouin A, Wilton DK, Funk K, DeMasters BK, Jiang X, Bowen JR, Mennerick S, Robinson JK, Garbow JR, Tyler KL, Suthar MS, Schmidt RE, Stevens B, Klein RS. A complement-microglial axis drives synapse loss during virus-induced memory impairment. Nature. 2016 Jun 23;534(7608):538–43. doi: 10.1038/nature18283.

Walsh MC, Choi Y. Biology of the RANKL-RANK-OPG System in Immunity, Bone, and Beyond. Front Immunol. 2014 Oct 20;5:511. doi: 10.3389/fimmu.2014.00511.

Walsh MC, Choi Y. Regulation of T cell-associated tissues and T cell activation by RANKL-RANK-OPG. J Bone Miner Metab. 2021 Jan;39(1):54–63. doi: 10.1007/s00774-020-01178-y.

Wanner R, Knöll B. Interference with SRF expression in skeletal muscles reduces peripheral nerve regeneration in mice. Sci Rep. 2020 Mar 24;10(1):5281. doi: 10.1038/s41598-020-62231-4.

Wang F, Flanagan J, Su N, Wang LC, Bui S, Nielson A, Wu X, Vo HT, Ma XJ, Luo Y. RNAscope: a novel in situ RNA analysis platform for formalin-fixed, paraffin-embedded tissues. J Mol Diagn. 2012 Jan;14(1):22–9. doi: 10.1016/j.jmoldx.2011.08.002.

Wang J, Campos B, Jamieson GA Jr, Kaetzel MA, Dedman JR. Functional elimination of calmodulin within the nucleus by targeted expression of an inhibitor peptide. J Biol Chem. 1995 Dec 22;270(51):30245–8. doi: 10.1074/jbc.270.51.30245.

Yamashita T, Vavladeli A, Pala A, Galan K, Crochet S, Petersen SSA, Petersen CCH. Diverse Long-Range Axonal Projections of Excitatory Layer 2/3 Neurons in Mouse Barrel Cortex. Front Neuroanat. 2018 May 1;12:33. doi: 10.3389/fnana.2018.00033.

Yan EB, Satgunaseelan L, Paul E, Bye N, Nguyen P, Agyapomaa D, Kossmann T, Rosenfeld JV, Morganti-Kossmann MC. Post-traumatic hypoxia is associated with prolonged cerebral cytokine production, higher serum biomarker levels, and poor outcome in patients with severe traumatic brain injury. J Neurotrauma. 2014 Apr 1;31(7):618–29. doi: 10.1089/neu.2013.3087.

Yan J, Bengtson CP, Buchthal B, Hagenston AM, Bading H. Coupling of NMDA receptors and TRPM4 guides discovery of unconventional neuroprotectants. Science. 2020 Oct 9;370(6513):eaay3302. doi: 10.1126/science.aay3302.

Yang X, Wang X, Chi M, Zhang M, Shan H, Zhang QH, Zhang J, Shi J, Zhang JZ, Wu RM, Li YL. Osteoprotegerin mediate RANK/RANKL signaling inhibition eases asthma inflammatory reaction by affecting the survival and function of dendritic cells. Allergol Immunopathol (Madr). 2019 Mar-Apr;47(2):179–184. doi: 10.1016/j.aller.2018.06.006.

Yu Y, Oberlaender K, Bengtson CP, Bading H. One nuclear calcium transient induced by a single burst of action potentials represents the minimum signal strength in activity-dependent transcription in hippocampal neurons. Cell Calcium. 2017 Jul;65:14–21. doi: 10.1016/j.ceca.2017.03.003.

Yun TJ, Tallquist MD, Aicher A, Rafferty KL, Marshall AJ, Moon JJ, Ewings ME, Mohaupt M, Herring SW, Clark EA. Osteoprotegerin, a crucial regulator of bone metabolism, also regulates B cell development and function. J Immunol. 2001 Feb 1;166(3):1482–91. doi: 10.4049/jimmunol.166.3.1482.

Zhang SJ, Buchthal B, Lau D, Hayer S, Dick O, Schwaninger M, Veltkamp R, Zou M, Weiss U, Bading H. A signaling cascade of nuclear calcium-CREB-ATF3 activated by synaptic NMDA receptors defines a gene repression module that protects against extrasynaptic NMDA receptor-induced neuronal cell death and ischemic brain damage. J Neurosci. 2011 Mar 30;31(13):4978–90. doi: 10.1523/JNEUROSCI.2672-10.2011.

Zhang SJ, Zou M, Lu L, Lau D, Ditzel DA, Delucinge-Vivier C, Aso Y, Descombes P, Bading H. Nuclear calcium signaling controls expression of a large gene pool: identification of a gene program for acquired neuroprotection induced by synaptic activity. PLoS Genet. 2009 Aug;5(8):e1000604. doi: 10.1371/journal.pgen.1000604.

Zhang Y, Williams PR, Jacobi A, Wang C, Goel A, Hirano AA, Brecha NC, Kerschensteiner D, He Z. Elevating Growth Factor Responsiveness and Axon Regeneration by Modulating Presynaptic Inputs. Neuron. 2019 Jul 3;103(1):39–51.e5. doi: 10.1016/j.neuron.2019.04.033.

