## supplementary table 3 for "Neuronal nuclear calcium signaling suppression of microglial reactivity is mediated by osteoprotegerin after traumatic brain injury"

| **Antibody** | **Host** | **Dilution** | **Company** | **Catalog Number** |
| --- | --- | --- | --- | --- |
| **NeuN** | Guinea Pig | 1 to 500 | SySy | 266004 |
| **NeuN (clone 60)** | Mouse | 1 to 300 | Millipore | MAB377 |
| **Phospho-CREB (Ser133) (87G3)** | Rabbit | 1 to 500 | CST | 9198S |
| **RFP** | Camel | 1 to 500 | Nanotech | N0404-AT565-L |
| **Iba1** | Guinea Pig | 1 to 500 | SySy | 234004 |
| **Iba1** | Rabbit | 1 to 250 | Wako | 019-19741 |
| **GFAP** | Chicken | 1 to 500 | Abcam | Ab4676 |
| **TMEM119 (28-3)** | Rabbit | 1 to 100 | Abcam | Ab209064 |
| **CD11c (3.9)** | Mouse | 1 to 150 | Abcam | Ab11029 |
| **CST7** | Rabbit | 1 to 100 | Biorbyt | Orb101860 |
| **CD169** | Rat | 1 to 200 | Bio-Rad | MCA947GA |
| **SHANK2** | Rabbit | 1 to 500 | Homemade | SA5192 |
| **SHANK3** | Rabbit | 1 to 500 | Homemade | Tier 2 |
| **Dapi** | - | 1 to 1000 | Thermo | 62247 |
| **Anti-Mouse Alexa Fluor 405** | Donkey | 1 to 500 | Abcam | Ab175658 |
| **Anti-Guinea Pig Alexa Fluor 405** | Goat | 1 to 500 | Abcam | Ab175678 |
| **Anti-Chicken CF488A** | Donkey | 1 to 500 | Biotium | 20166 |
| **Anti-Rabbit Alexa Fluor 488** | Donkey | 1 to 500 | Invitrogen | A21206 |
| **Anti-Guinea Pig Alexa Fluor 488** | Goat | 1 to 500 | Invitrogen | A11073 |
| **Anti-Mouse Alexa Fluor 568** | Goat | 1 to 500 | Invitrogen | A11031 |
| **Anti-Guinea Pig CF633** | Donkey | 1 to 500 | Biotium | 20171 |
| **Anti-Rat Alexa Fluor 647** | Chicken | 1 to 500 | Invitrogen | A21472 |
| **Anti-Mouse Alexa Fluor 647** | Donkey | 1 to 500 | Invitrogen | A31571 |

**Antibodies**

**Constructs**

| **Construct** | **Source** | **Basepairs** |
| --- | --- | --- |
| P5E18VD2/9 | Heidelberg (AG Kiel-Müller) | 7329 |
| pAd-DeltaF6 | Addgene 112867 | 15420 |
| hSyn-mCherry.NLS.myc | AG Bading | 5765 |
| hSyn-PV.NLS.mCherry | AG Bading | 6049 |
| hSyn-CaMBP4.flag.mCherry | AG Bading | 5953 |
| hSyn-TNFRSF11b-P2A-mCherry | Vector Biosciences | 6571 |
| CAG-A7-floxed-PSAM(L141F,Y115F)-GlyR-GFP-WPRE | Addgene 32481 | 7454 |
