## Supplementary figures and images for "Neuronal nuclear calcium signaling suppression of microglial reactivity is mediated by osteoprotegerin after traumatic brain injury"

### supplementary figure 1

Fröhlich et al., Supplementary Figure 1.pdf
2106 x 1282

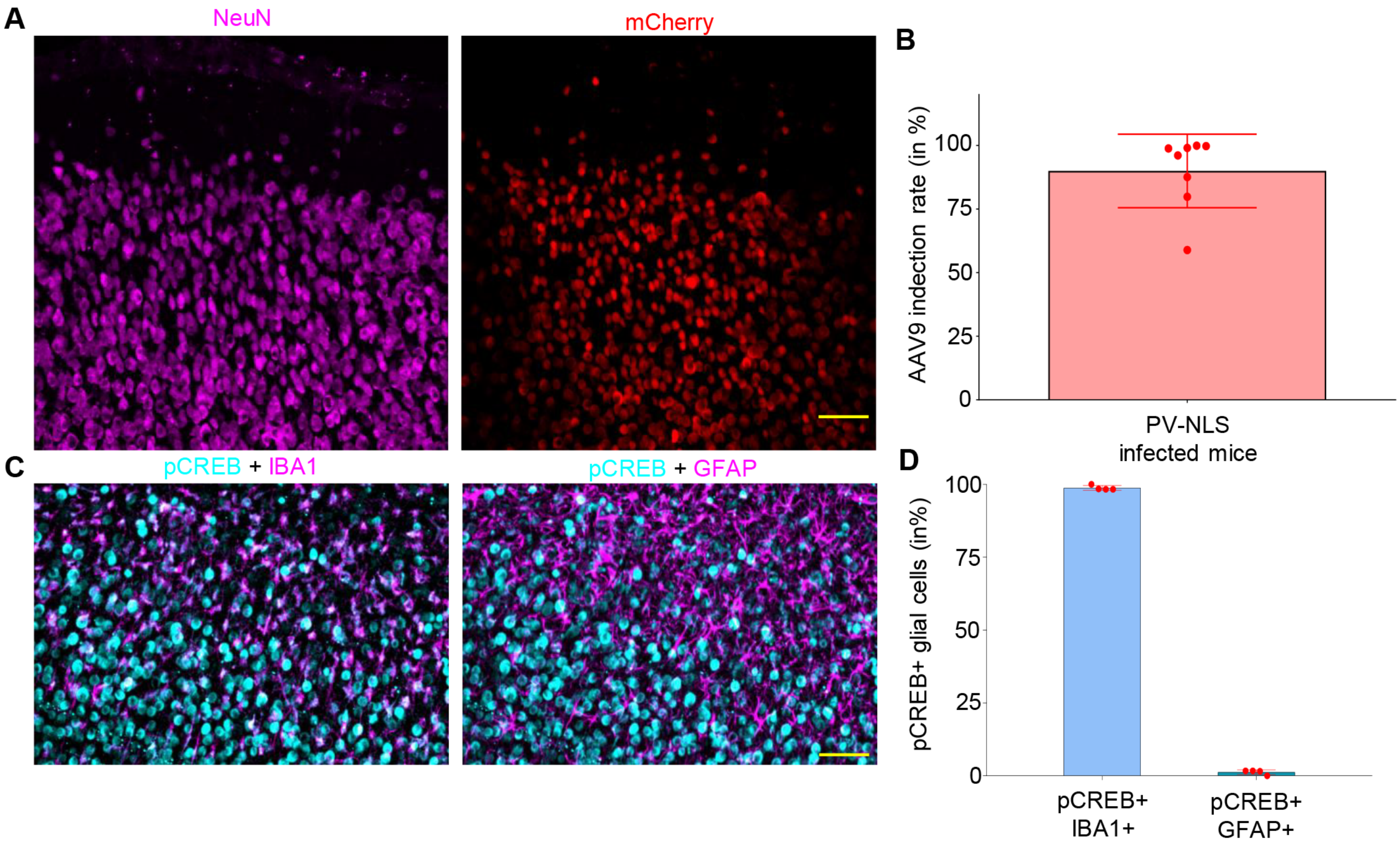

### supplementary figure 2

Fröhlich et al., Supplementary Figure 2.pdf
1962 x 1606

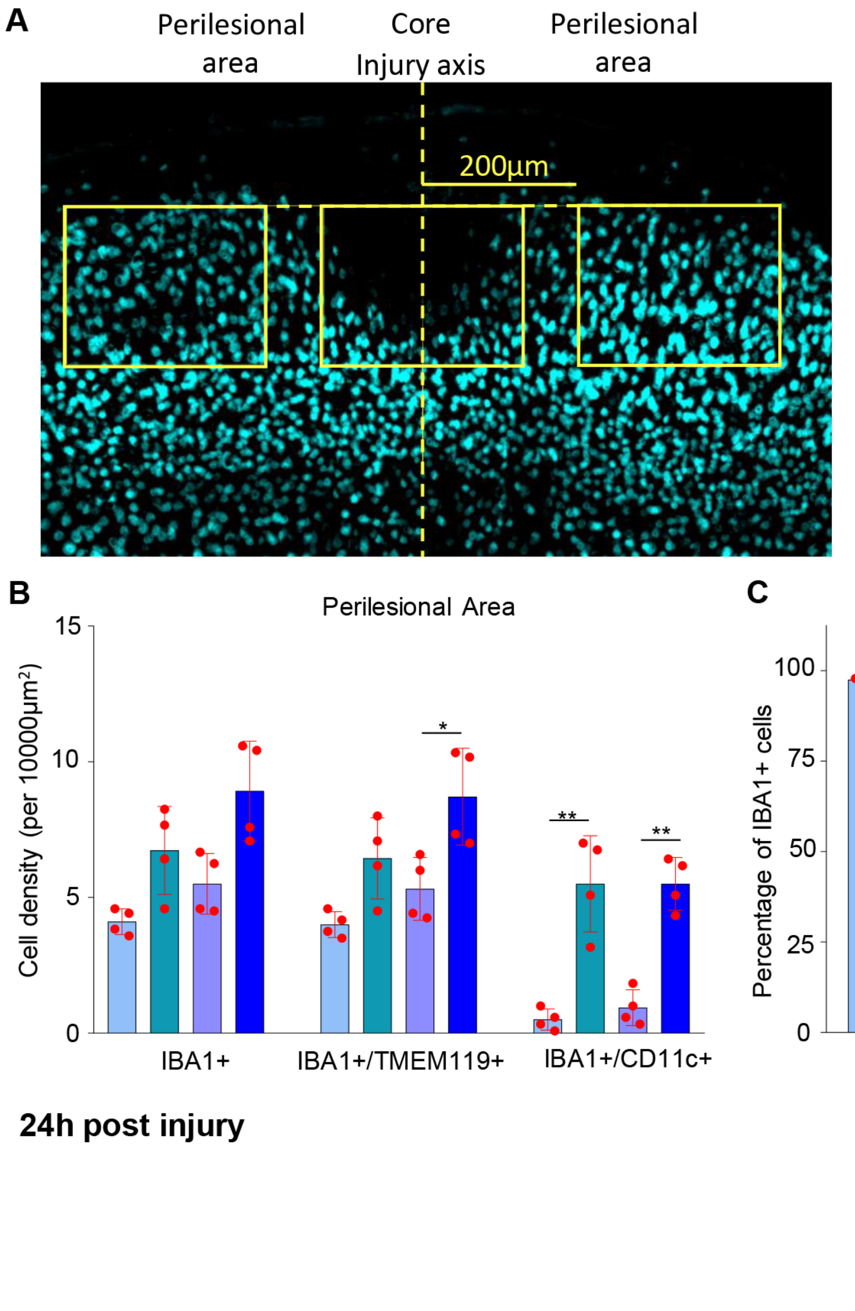


**24h post injury**

### supplementary figure 3

Fröhlich et al., Supplementary Figure 3.pdf

2046 x 1042

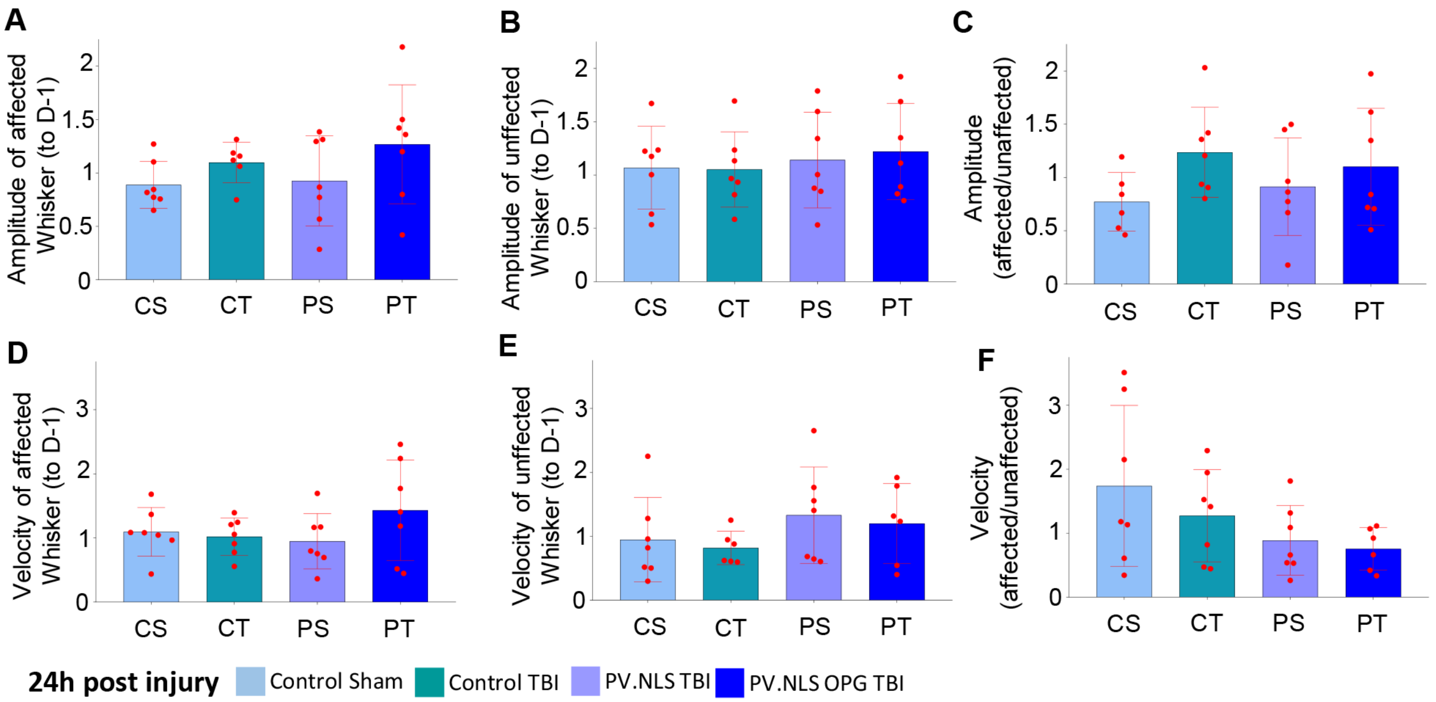

### supplementary figure 4

Fröhlich et al., Supplementary Figure 4.pdf
1867 x 1030

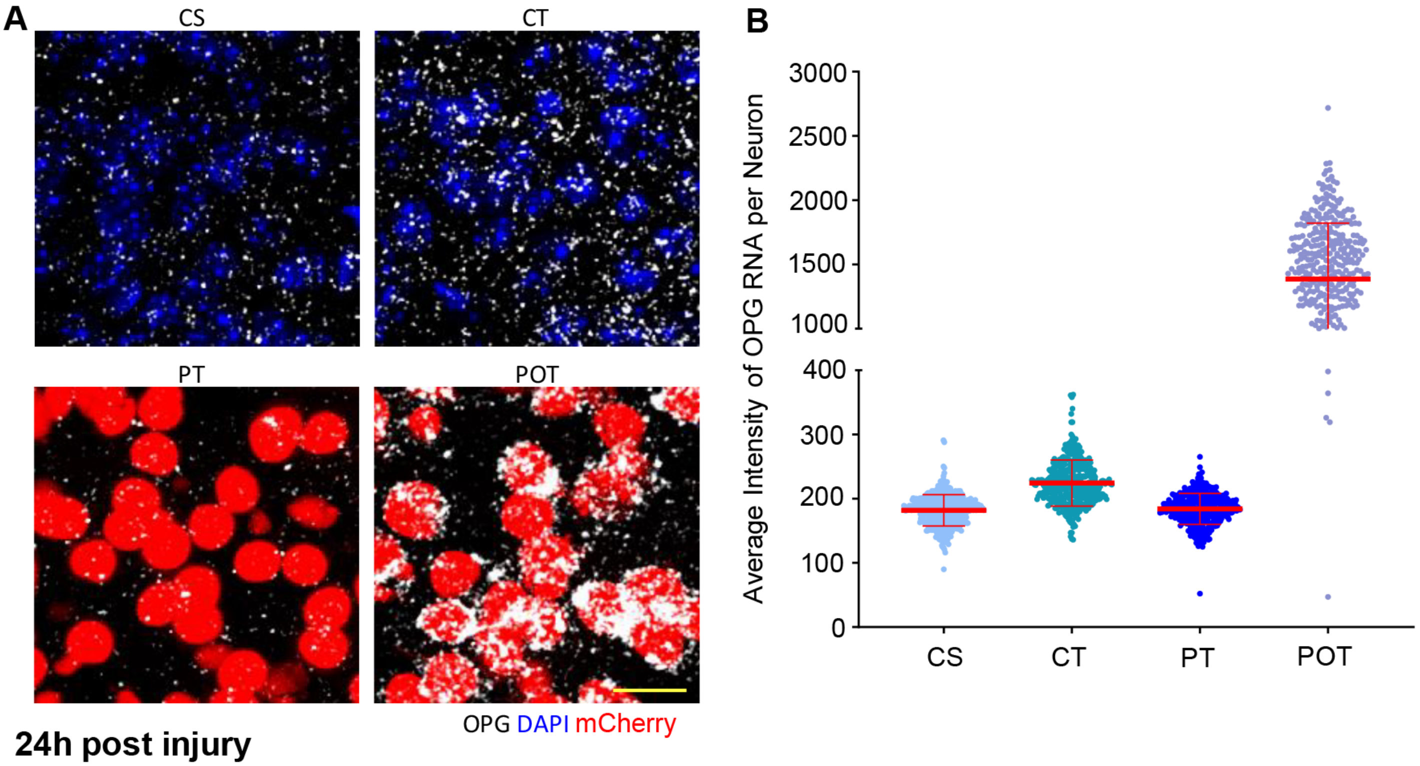


**24h post injury**
